# Almost Free Enhancement of Multi-Population PRS: From Data-Fission to Pseudo-GWAS Subsampling

**DOI:** 10.1101/2025.06.16.659952

**Authors:** Leqi Xu, Yikai Dong, Xiaowei Zeng, Zeyu Bian, Geyu Zhou, Leying Guan, Hongyu Zhao

## Abstract

Many multi-population polygenic risk score (PRS) methods have been proposed to improve prediction accuracy in underrepresented populations; however, no single method outperforms other methods across all data scenarios. Although integrating PRS results across multiple methods and populations may lead to more accurate predictions, this approach may be limited by the availability of individual-level tuning data to calculate combination weights. In this manuscript, we introduce MIXPRS, a robust PRS integration framework based on data fission principles, to effectively combine multiple multi-population PRS methods using only genome-wide association study (GWAS) summary statistics from multiple populations. Specifically, MIXPRS employs SNP pruning to mitigate linkage disequilibrium (LD) mismatch between the training GWAS summary statistics and LD reference panels, and utilizes non-negative least squares regression to robustly estimate PRS combination weights. Extensive simulations and real-data analyses involving 22 continuous traits and four binary traits across five populations from the UK Biobank and All of Us datasets demonstrate that MIXPRS consistently outperforms the existing methods in prediction accuracy. Because MIXPRS relies solely on GWAS summary statistics, it enjoys broad accessibility, robustness, and generalizability for underrepresented populations.

## Introduction

Historically, genetic analyses have predominantly focused on European populations due to the availability of large European cohorts [1, 2]. It has been found that the genetic findings may not be generalized to other populations [3, 4]. To bridge this gap, there has been an increasing emphasis on generalizing genetic analyses beyond European populations. There has been an increase of non-European genome-wide association studies (GWAS) [5, 6, 7, 8, 9, 10, 11, 12, 13] coupled with the development of multi-population polygenic risk score (PRS) methods [14, 15, 16, 17, 18, 19, 20, 21, 22, 23, 24, 25, 26, 27]. These efforts have substantially enhanced diversity and improved the accuracy and generalizability of genetic risk predictions across diverse populations.

Despite these advancements, no multi-population PRS method consistently demonstrates superior performance across diverse populations and traits [14, 19, 20, 21]. This inconsistency arises primarily from differing underlying methods assumptions and the varying genetic architecture among populations and traits. Consequently, integrating multiple PRS methods is essential for robust and consistent predictive performance across populations and traits.

However, PRS methods integration typically requires individual-level tuning data, which are often unavailable, particularly for underrepresented populations. Therefore, it is important to develop PRS integration methods relying solely on GWAS summary statistics.

Moreover, current GWAS analyses often use meta-analysis to combine multiple cohorts, increasing statistical power but yielding a single, aggregated GWAS summary statistics dataset [5, 6, 7, 8, 9, 10, 11, 12, 13]. This aggregation precludes access to independent GWAS datasets necessary for separately training PRS models and effectively combining across PRS methods, substantially increasing overfitting risks.

To address this challenge, “data fission” [28, 29, 30] has recently been introduced that partitions a single dataset into two independent subsets, preserving genuine signals while introducing independent residual variations. Originally developed outside the GWAS context, data fission has been effectively adapted into pseudo-GWAS subsampling [31, 32, 33]. This adaptation enables researchers to derive independent training and tuning GWAS datasets from a single GWAS summary statistics, mimicking independent GWAS cohorts.

Moreover, PRS integration can be reformulated to operate entirely on GWAS summary statistics by transforming individual-level optimization problems into equivalent summary-level formulations. As a result, the PRS integration framework can both train individual PRS models and combine across methods using only a single GWAS summary statistics dataset.

Despite this progress, current PRS integration methods primarily focus on single-populations, which are less effective for underrepresented populations [31, 32, 33]. Furthermore, the existing PRS integration methods often neglect linkage disequilibrium (LD) mismatch between the training GWAS summary statistics and LD reference panels, further exacerbating overfitting risks [31, 32, 33].

To overcome these limitations, we introduce a multi-population PRS integration framework (MIXPRS), specifically designed to combine multiple multi-population PRS methods using only GWAS summary statistics. MIXPRS applies SNP pruning to select a subset of approximately independent SNPs, thereby mitigating LD mismatch, and employs non-negative least squares regression (NNLS) to obtain robust and stable estimates of PRS combination weights.

To demonstrate the practical advantage and better prediction accuracy of MIXPRS, we conducted extensive simulations and real-data analyses, benchmarking against seven established multi-population PRS methods. Specifically, we considered 22 continuous and four binary traits evaluated across five distinct populations (European (EUR), East Asian (EAS), African (AFR), South Asian (SAS), and Admixed American (AMR)) using the UK Biobank (UKBB) [34] and All of Us (AoU) datasets [35]. By integrating the strengths of various multi-population PRS methods, MIXPRS consistently achieves better prediction accuracy across diverse data scenarios. Crucially, MIXPRS requires only GWAS summary statistics, eliminating the dependency on individual-level tuning data. This significantly enhances the robustness and generalizability of genetic risk predictions across diverse populations.

## Results

### Overview of MIXPRS

MIXPRS introduces a PRS integration framework based on the data fission principle, effectively integrating PRS derived from multiple populations and methods using only GWAS summary statistics. The MIXPRS framework consists of three key steps (Figure 1): (1) GWAS subsampling in MIXPRS to create independent subsampled training and tuning GWAS datasets; (2) Estimation of MIXPRS combination weights for different PRS methods; and (3) Derivation of the final integrated MIXPRS. The detailed implementations of MIXPRS are provided in the Methods section.

**Figure 1.**
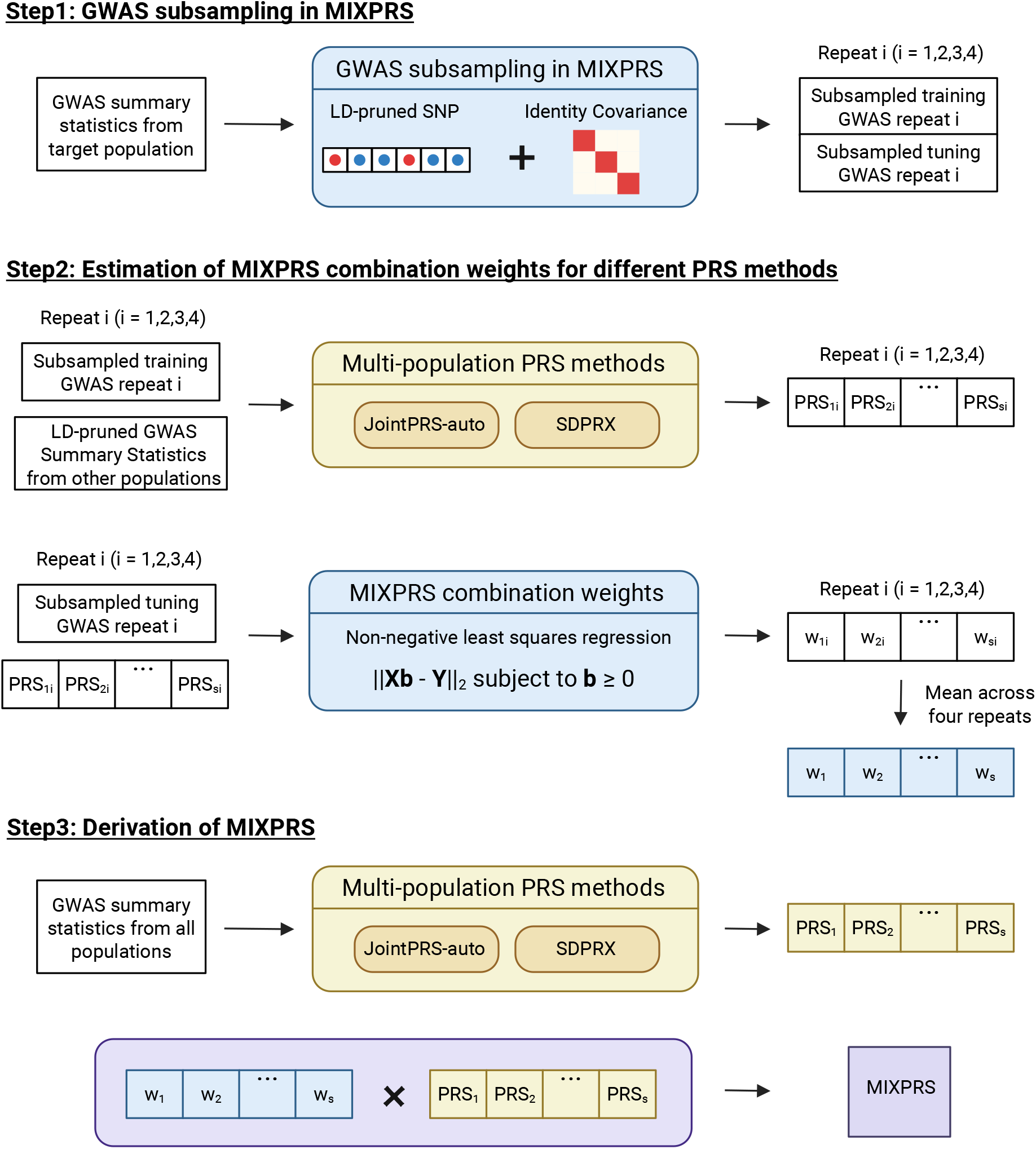
MIXPRS workflow. MIXPRS integrates multi-population PRS methods through three key steps: (1) Step1: GWAS subsampling in MIXPRS: MIXPRS generates subsampled training and tuning GWAS datasets from the original GWAS summary statistics of the target population, using only LD-pruned SNPs with an identity covariance structure. (2) Step2: Estimation of MIXPRS combination weights for different PRS methods: This step consists of two sub-steps: (i) Calculation of PRS using subsampled training GWAS along with LD-pruned GWAS summary statistics from other populations via JointPRS-auto and SDPRX. (ii) Estimation of PRS combination weights by applying NNLS to the subsampled tuning GWAS and previously calculated PRS. (3) Step3: Derivation of MIXPRS: PRS are computed using the original GWAS summary statistics from all populations via JointPRS-auto and SDPRX, then combined using the PRS combination weights obtained in Step 2 to produce the final MIXPRS. This figure was created in BioRender.

#### Step1: GWAS subsampling in MIXPRS

This initial step utilizes GWAS summary statistics from the target population. The GWAS subsampling method in MIXPRS leverages data fission to partition a single original GWAS dataset into independent subsampled training and tuning GWAS datasets [28, 29]. To mitigate the effects of LD mismatch between the training GWAS summary statistics and LD reference panels, MIXPRS utilizes LD-pruned SNPs (retaining SNPs with pairwise correlation below 0.5 within 250 kb sliding windows) [36] and adopts an identity covariance structure.

#### Step2: Estimation of MIXPRS combination weights for different PRS methods

This step involves two sub-steps. First, the subsampled training GWAS datasets from the target population, combined with LD-pruned GWAS summary statistics from other populations, are input into established multi-population PRS methods (JointPRS-auto [14] and SDPRX [18]) to compute LD-pruned PRS beta values. Second, weights for combining these PRS methods, each leveraging the computed LD-pruned PRS beta values, are estimated via non-negative least squares regression (NNLS) [37], utilizing the subsampled tuning GWAS datasets. The NNLS approach enhances the robustness and accuracy in estimating PRS combining weights.

#### Step3: Derivation of MIXPRS

In the final step, the original GWAS summary statistics from all populations are utilized to calculate full SNPs PRS beta values using JointPRS-auto and SDPRX. MIXPRS subsequently integrates these full SNPs PRS beta values using the weights derived from Step2, resulting in the final integrated MIXPRS, which offers improved predictive performance across multiple populations.

### Simulation results

We conducted comprehensive simulations to evaluate the performance of MIXPRS compared to seven existing multi-population PRS methods: JointPRS [14], XPASS [15], SDPRX [18], PRS-CSx [17], MUSSEL [20], PROSPER [21], and BridgePRS [22]. JointPRS, PRS-CSx, MUSSEL, and PROSPER utilized GWAS summary statistics from all available populations (EUR, EAS, AFR, SAS, and AMR). In contrast, XPASS and SDPRX incorporated GWAS summary statistics exclusively from two populations—the European population and the specific non-European target population.

As shown in Figure 2, Table 1 and Table S5, MIXPRS consistently outperformed other PRS methods across all causal SNP proportions (*p* = 0.1, 0.01, 0.001, 5 *×* 10^−4^) and both non-European training sample sizes (*N* = 25, 000 and 90, 000) for the four non-European populations (EAS, AFR, SAS, and AMR). Specifically, the average improvements of MIXPRS compared to other methods across five repeats and four causal SNP proportions ranged from 4.29% to 146.48% at *N* = 25, 000 and from 6.93% to 114.73% at *N* = 90, 000 for the EAS population, from 5.41% to 155.92% at *N* = 25, 000 and from 8.31% to 156.11% at *N* = 90, 000 for the AFR population, from 4.03% to 136.94% at *N* = 25, 000 and 5.69% to 90.92% at *N* = 90, 000 for the SAS population, and from 4.51% to 164.55% at *N* = 25, 000 and from 5.64% to 173.55% for the AMR population. These results demonstrate the advantage of using MIXPRS to integrate PRS across multiple populations and methods.

**Table 1.** Average improvement of MIXPRS over each method in simulations.

| Pop | Training sample | JointPRS | XPASS | SDPRX | PRS-CSx | MUSSEL | PROSPER | BridgePRS |
| --- | --- | --- | --- | --- | --- | --- | --- | --- |
| EAS | 25K | 4.29% | 146.48% | 7.27% | 10.12% | 28.41% | 32.53% | 73.99% |
|  | 90K | 6.93% | 57.03% | 7.92% | 37.82% | 60.61% | 67.2% | 114.73% |
| AFR | 25K | 5.41% | 155.92% | 7.46% | 11.51% | 20.85% | 54.51% | 96.05% |
|  | 90K | 8.31% | 78.26% | 10.25% | 49.51% | 66.27% | 114% | 156.11% |
| SAS | 25K | 4.03% | 136.94% | 8.58% | 8.19% | 43.55% | 26.96% | 56.17% |
|  | 90K | 5.69% | 46.71% | 9.81% | 33.2% | 75.54% | 57.22% | 90.92% |
| AMR | 25K | 4.51% | 164.55% | 7.22% | 8.63% | 34.58% | 35.97% | 53.84% |
|  | 90K | 5.64% | 173.55% | 8.81% | 31.9% | 64.08% | 66.99% | 85.35% |

**Figure 2.**
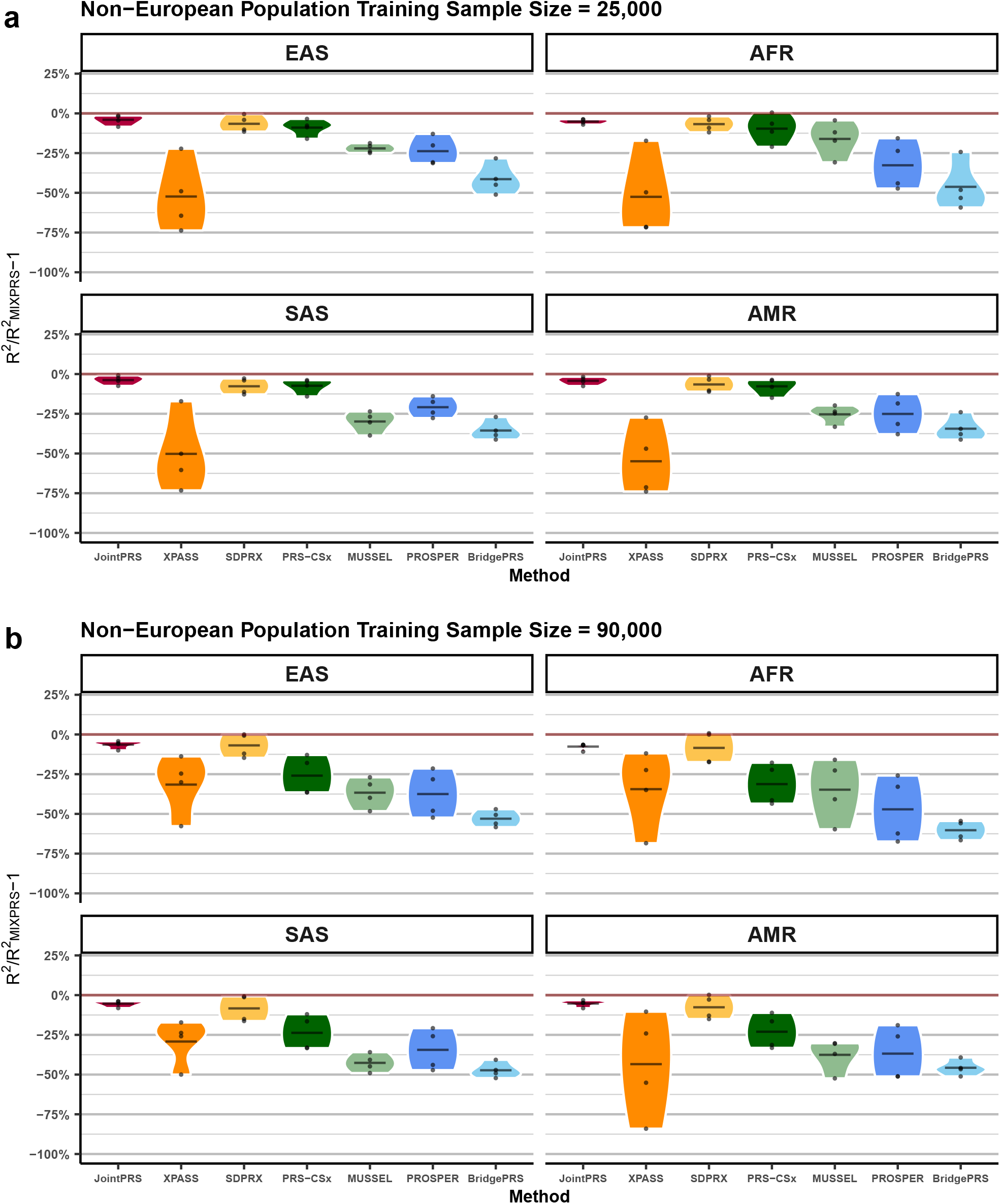
Relative performance of multi-population PRS methods compared to MIXPRS across varying sample sizes and genetic architectures in simulations. **a–b**, Simulations were performed with total heritability fixed at *h*^2^ = 0.4, cross-population genetic correlation fixed at *ρ* = 0.8, and four causal SNP proportions: *p* = 0.1, 0.01, 0.001, 5 *×* 10^−4^. Non-European populations (EAS, AFR, SAS, and AMR) have training sample sizes of **a** *N*_train_ = 90, 000 and **b** *N*_train_ = 25, 000, while the EUR population size is fixed at *N*_train_ = 311, 600. Each dot represents the mean relative performance, defined as 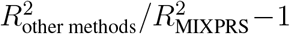 across five simulation replicates for each scenario.

Further comparisons were conducted between MIXPRS and its component methods (JointPRS-auto and SDPRX) using a GWAS training sample size of 100, 000 individuals, noting that none of these methods require individual-level tuning data. Figure S1 and Table S6 illustrate that JointPRS-auto demonstrated superior performance in scenarios with higher causal SNP proportions, whereas SDPRX excelled in scenarios involving lower causal SNP proportions. Importantly, MIXPRS consistently achieved performance equal to or better than both component methods across all scenarios, effectively combining their strengths.

Finally, we assessed the effectiveness of the SNP pruning strategy within the GWAS subsampling step of MIXPRS to address the LD mismatch issue. When there is LD mismatch between the simulated data and the LD reference panel, residuals from the subsampled training and tuning GWAS datasets become correlated, which can lead to overfitting and reduced prediction accuracy. To evaluate this, we analyzed residual correlations and compared prediction accuracy across various GWAS subsampling strategies (details are provided in the Methods section). Figure S2a and Table S7 demonstrate that SNP pruning significantly reduces residual correlations compared to utilizing all available SNPs, effectively mitigating overfitting due to LD mismatch. Notably, using an identity covariance structure combined with SNP pruning resulted in slightly negative residual correlations for four non-European populations. Lower residual correlations were particularly evident in non-European populations, reflecting closer LD alignment between simulated genetic data and the LD reference panel. In contrast, European populations from the UKBB exhibited higher residual correlations, indicating greater real-world LD complexity.

Figure S2b and Table S8 illustrate that MIXPRS employing the SNP pruning strategy outperformed approaches using the full SNP set, specifically under scenarios involving sparse causal SNP proportions. These results underscore the critical role of SNP pruning in mitigating LD mismatch and enhancing prediction accuracy within the MIXPRS framework. However, the optimal choice of pruning strategy varied by population. For European populations, which closely align with real-data analyses, using the SNP pruning strategy combined with an identity covariance structure yielded the greatest improvement (2.34%) compared to the full SNP set with a LD reference panel across four causal SNP proportions, whereas using the SNP pruning strategy with the LD reference panel provided a minor improvement (0.09%). Conversely, for non-European populations, due to the similarity between the simulated genetic data and the LD reference panel structure, pruning SNPs combined with an LD reference panel strategy provided modest benefits (no improvement for EAS, 0.95% for AFR, 0.54% for SAS, and 0.94% for AMR), while the identity covariance strategy offered no improvement across the four non-European populations.

### Effectiveness and Robustness of MIXPRS

We evaluated the effectiveness and robustness of MIXPRS through multiple comparative analyses across 22 continuous and four binary traits in UKBB for the four non-European populations (EAS, AFR, SAS, and AMR). As illustrated in Figure 3 and Table S9, we initially compared MIXPRS (Prune NNLS), which uses pruned SNPs with an identity covariance matrix in the GWAS subsampling step combined with NNLS in the PRS combining weights estimation step, to an alternative approach (Full Linear) using full SNP sets with the LD reference panel based on the 1000 Genomes Project [38] in the GWAS subsampling step combined with linear regression in the PRS combining weights estimation step. Prune NNLS showed substantial improvements in comparison to Full Linear: 9.48% across 22 continuous and four binary traits in the EAS population, 4.49% across nine continuous and two binary traits in the AFR population, 33.33% across four continuous traits in the SAS population, and 43.52% across four continuous traits in the AMR population.

**Figure 3.**
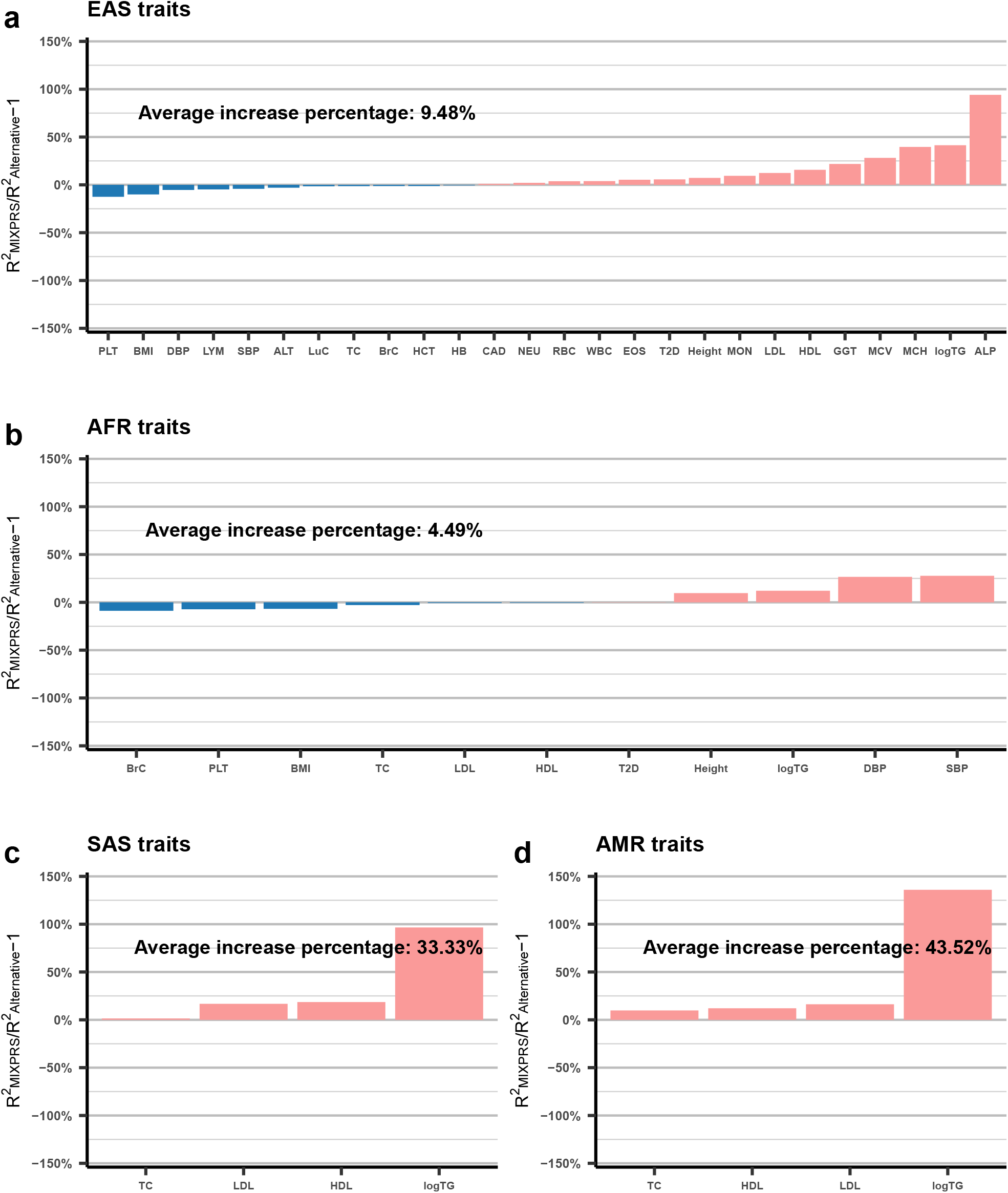
Relative improvement of MIXPRS over the alternative method across 26 traits in UKBB without tuning data. **a–d**, Relative improvement of MIXPRS compared to the alternative method evaluated across 26 traits in four non-European populations (**a** EAS; **b** AFR; **c** SAS; and **d** AMR) in UKBB, without using individual-level tuning data. Relative performance was calculated as 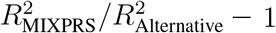 for quantitative traits and AUC_MIXPRS_*/*AUC_Alternative_ − 1 for binary traits. Pink bars indicate positive improvement (MIXPRS outperforms the alternative method), while blue bars indicate negative improvement (the alternative method outperforms MIXPRS).

To separately assess the contributions of the SNP pruning strategy and the NNLS approach, we performed additional comparisons. First, pruned SNPs combined with linear regression (Prune Linear) were compared to Full Linear. As shown in Figure S3 and Table S9, SNP pruning alone improved performance by 3.57% in EAS, 1.85% in AFR, 27.17% in SAS, and 28.51% in AMR populations. Second, Prune NNLS was compared to Prune Linear, showing additional performance gains of 5.73% in EAS, 2.59% in AFR, 4.57% in SAS, and 11.47% in AMR populations by using NNLS (Figure S4 and Table S9). These analyses demonstrate significant and independent benefits from both the SNP pruning strategy and the NNLS approach within MIX-PRS.

We further assessed the robustness of MIXPRS concerning the choice of SNPs covariance structure and SNP pruning lists in real-data analyses. Figure S5 and Table S10 demonstrate robust MIXPRS performance when employing either an identity covariance matrix or an LD reference matrix, with slightly better overall results observed using the identity covariance matrix across all populations. This finding contrasts with simulation results, where non-European populations performed better using SNP pruning combined with the LD reference panel. This discrepancy occurs because, in simulations, non-European genetic data were simulated and closely matched the LD reference panel structure. In contrast, the European population simulations used the observed genetic data from UKBB, and showed greater performance benefits with SNP pruning combined with an identity covariance structure, aligning closely with our real-data findings. Therefore, due to significant LD mismatch inherent in real genetic data, employing the identity covariance structure is recommended in practice.

Additionally, Figure S6 and Table S11 illustrate robustness in MIXPRS performance across different SNP pruning lists. By default, MIXPRS prioritizes variants with higher non-major allele frequencies (snplist 1), with subsequent lists (snplist 2, snplist 3, snplist 4) prioritizing remaining SNPs. Among these, snplist 1 yielded slightly better predictive accuracy.

### MIXPRS performance benchmarking without tuning data (UKBB)

To evaluate the prediction accuracy of MIXPRS, we first benchmarked MIXPRS in the scenario when no individual-level tuning data is available. Specifically, we evaluated prediction accuracy across 22 continuous and four binary traits in UKBB in four non-European populations (EAS, AFR, SAS, and AMR). MIXPRS was compared to four existing multi-population PRS methods applicable to this scenario: JointPRS-auto, SDPRX, PRS-CSx-auto, and XPASS.

As shown in Figure 4, Table 2, Table S12, and Table S13, MIXPRS consistently improved predictive performance compared to existing methods across multiple populations when no individual-level tuning data is available. Specifically, average improvements across traits ranged from 5.17% to 54.18% in the EAS population, from 14.29% to 79.01% in the AFR population, from 16.25% to 103.30% in the SAS population, and from 15.62% to 116.52% in the AMR population. These consistent improvements highlight the effectiveness and robustness of MIXPRS across diverse populations.

**Table 2.**
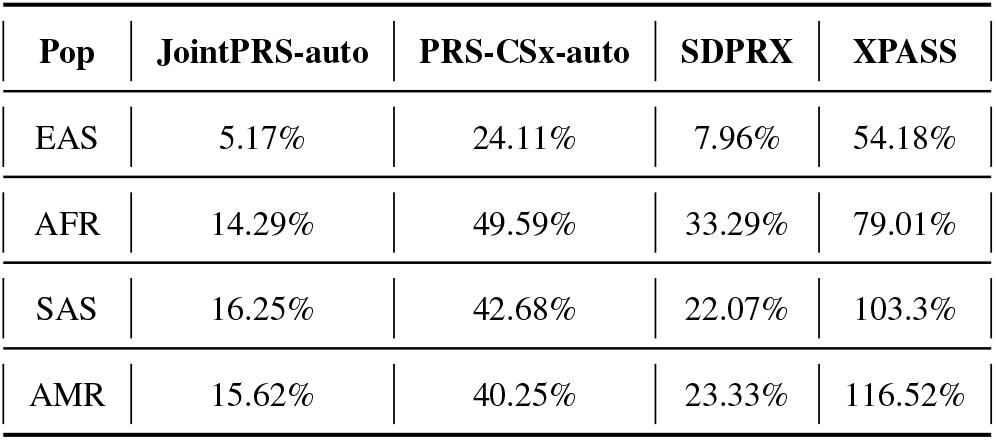
Average improvement of MIXPRS over each method without tuning data (UKBB).

| Pop | JointPRS-auto | PRS-CSx-auto | SDPRX | XPASS |
| --- | --- | --- | --- | --- |
| EAS | 5.17% | 24.11% | 7.96% | 54.18% |
| AFR | 14.29% | 49.59% | 33.29% | 79.01% |
| SAS | 16.25% | 42.68% | 22.07% | 103.3% |
| AMR | 15.62% | 40.25% | 23.33% | 116.52% |

**Figure 4.**
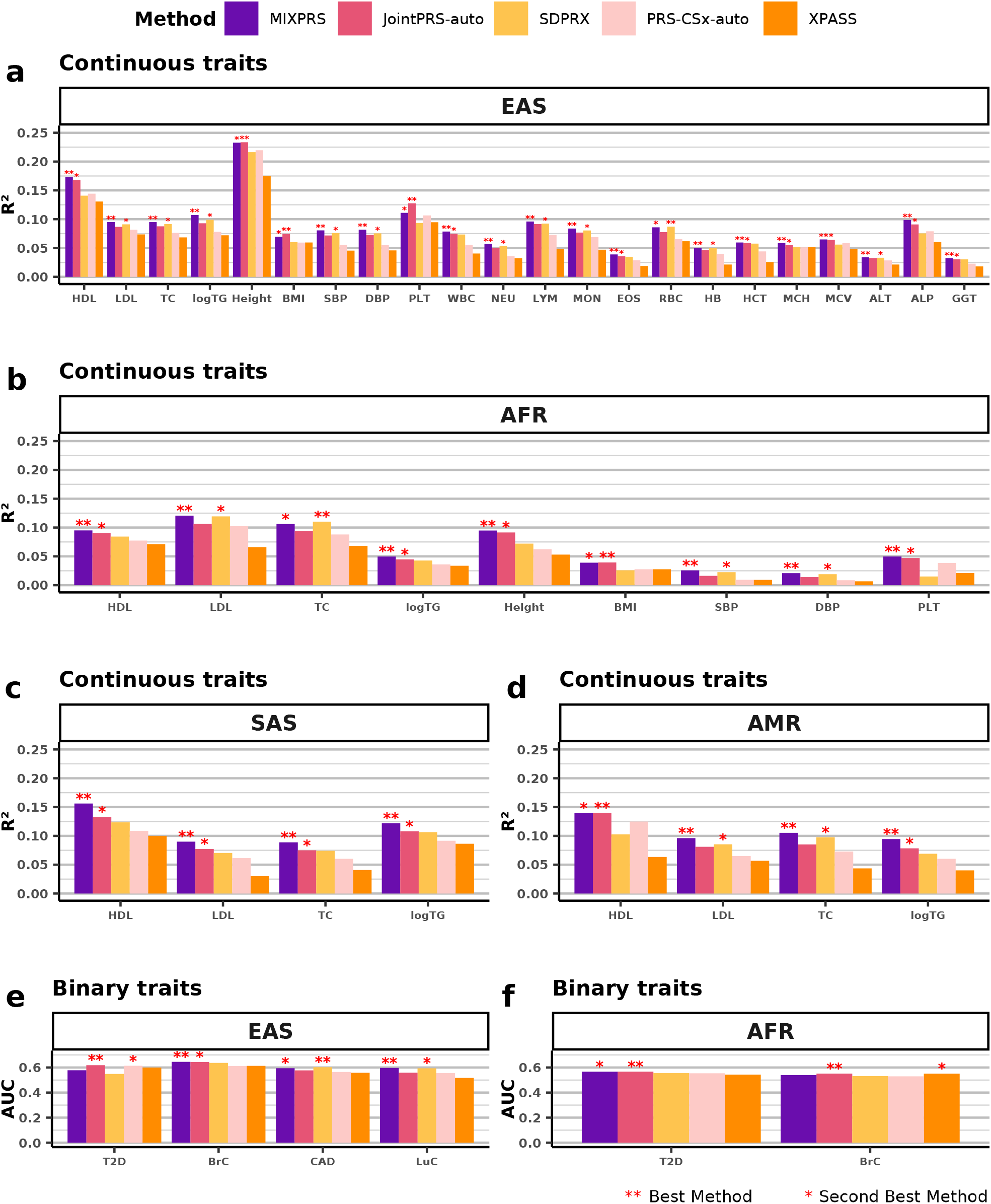
Prediction accuracy of multi-population PRS methods across 26 traits in UKBB without tuning data. **a–f**, Prediction accuracy of five multi-population PRS methods were evaluated across 26 traits in four non-European populations in UKBB without individual-level tuning data. Panels depict results for continuous traits in **a** EAS, **b** AFR, **c** SAS, and **d** AMR, and binary traits in **e** EAS and **f** AFR. Evaluation metrics were *R*^2^ for quantitative traits and AUC for binary traits, with results presented as bar plots. The best-performing and second-best-performing methods are indicated by two stars and one star, respectively, above the corresponding bars.

We further investigated MIXPRS performance relative to its component methods (JointPRS-auto and SDPRX) to better understand the effectiveness of combining PRS across methods and populations. As shown in Figure S7, Table S12, and Table S13, MIXPRS achieved the best performance for 18 out of 22 continuous traits and two out of four binary traits in the EAS population, seven out of nine continuous traits but none of the two binary traits in the AFR population, all four continuous traits in the SAS population, and three out of four continuous traits in the AMR population. These results demonstrate MIXPRS’s effectiveness in integrating the strengths of its component methods, leading to consistently improved prediction accuracy across multiple populations.

Moreover, to highlight the advantages of a multi-population PRS integration framework, we compared MIXPRS with PUMAS-EN, a recently developed single-population PRS integration framework capable of integrating across populations and methods [33]. For a fair comparison, we evaluated two versions of PUMAS-EN that differ in their use of LD reference panels and the number of integrated PRS methods (see Methods for details). The first version, PUMAS-EN, applied the PUMAS-EN framework using only the 1000 Genomes Project LD reference panel [38] throughout all stages, including GWAS subsampling, PRS training, and ensemble learning, matching the MIXPRS setup. This version integrated four commonly used PRS methods: lassosum [39], LDpred2[40], PRS-CS [41], and SBLUP [42]. The second version, PUMASEN paper, followed the setup described in the original publication [33], using the 1000 Genomes LD reference panel only for GWAS subsampling and UKBB LD reference panel for PRS training and ensemble learning. It incorporated nine PRS methods, including lassosum, LDpred2, PRS-CS, MegaPRS[43], SBayesR [44], DBSLMM [45], Vilma[46], and SBLUP.

As illustrated in Figure S8 and Table S14, MIXPRS consistently outperformed both PUMASEN versions across four lipid traits in three non-European populations (EAS, AFR, SAS). The AMR population was excluded to remain consistent with the original benchmarking approach used in the PUMAS-EN study. Specifically, MIXPRS achieved average performance improvements over PUMAS-EN and PUMAS-EN paper, respectively, of 300.97% and 37.55% in EAS, 101.68% and 14.21% in AFR, and 210.08% and 43.69% in SAS.

Notably, the substantial performance gap between PUMAS-EN and PUMAS-EN paper likely results from sensitivity to the included single-population PRS methods and the potential LD mismatch and associated overfitting when employing a single LD reference panel. MIXPRS effectively mitigates this issue via SNP pruning while still requiring only one LD reference panel and utilizing only two multi-population PRS methods. Furthermore, the marked improvement of MIXPRS over PUMAS-EN paper underscores the strength of multi-population PRS methods that jointly model genetic effects across populations. This joint modeling approach enhances predictive accuracy in non-European populations by borrowing genetic information from multiple populations, an advantage that cannot be matched by simply combining single-population PRS methods.

### MIXPRS performance benchmarking with tuning data (UKBB and AoU)

We also benchmarked the predictive performance of MIXPRS in the scenario where individual-level tuning data are available. We considered two scenarios: tuning and testing within the same cohort, and tuning and testing across different cohorts. For the same-cohort scenario, we performed five-fold cross-validation in the UKBB across 22 continuous and four binary traits in four non-European populations (EUR, EAS, AFR, and AMR). For the cross-cohort scenario, tuning was conducted in the UKBB and testing in AoU across nine continuous traits in two non-European populations (AFR and AMR). In both scenarios, we compared MIXPRS performance to seven existing methods: JointPRS, XPASS, SDPRX, PRS-CSx, MUSSEL, PROSPER, and BridgePRS.

As shown in Figure 5, Table 3, Table S15, and Table S16, MIXPRS consistently improved predictive performance compared to existing methods across multiple populations for the same-cohort scenario. Specifically, MIXPRS achieved average improvements across traits ranging from 1.70% to 642.68% for the EAS population, 0.70% to 147.62% for the AFR population, 0.19% to 106.17% for the SAS population, and 5.98% to 88.54% for the AMR population. In the cross-cohort scenario, as shown in Figure 6, Table 4, and Table S17, MIXPRS again demonstrated consistent improvements across populations, with average performance gains across traits ranging from 0.11% to 105.33% for the AFR population and from 5.84% to 185.14% for the AMR population. These results highlight the robustness and effectiveness of MIXPRS in enhancing predictive accuracy across diverse populations and evaluation scenarios.

**Table 3.** Average improvement of MIXPRS over each method when tuning and testing data are from the same cohort (UKBB).

| Pop | JointPRS | XPASS | SDPRX | PRS-CSx | MUSSEL | PROSPER | BridgePRS |
| --- | --- | --- | --- | --- | --- | --- | --- |
| EAS | 1.7% | 57.24% | 7.5% | 3.93% | 18.99% | 5.33% | 642.68% |
| AFR | 0.7% | 88.61% | 24.49% | 17.36% | 46.27% | 7.84% | 147.62% |
| SAS | 4.55% | 106.17% | 21.75% | 7.07% | 10.39% | 0.19% | 35.97% |
| AMR | 5.98% | 88.54% | 20.32% | 29.85% | 19.78% | 21.35% | 64.43% |

**Table 4.** Average improvement of MIXPRS over each method when tuning and testing data are from different cohorts (UKBB and AoU).

| Pop | JointPRS | XPASS | SDPRX | PRS-CSx | MUSSEL | PROSPER | BridgePRS |
| --- | --- | --- | --- | --- | --- | --- | --- |
| AFR | 0.11% | 54.69% | 25.72% | 7.4% | 16.72% | 7.73% | 105.33% |
| AMR | 5.84% | 185.14% | 24.73% | 17.18% | 86.98% | 12.45% | 67.52% |

**Figure 5.**
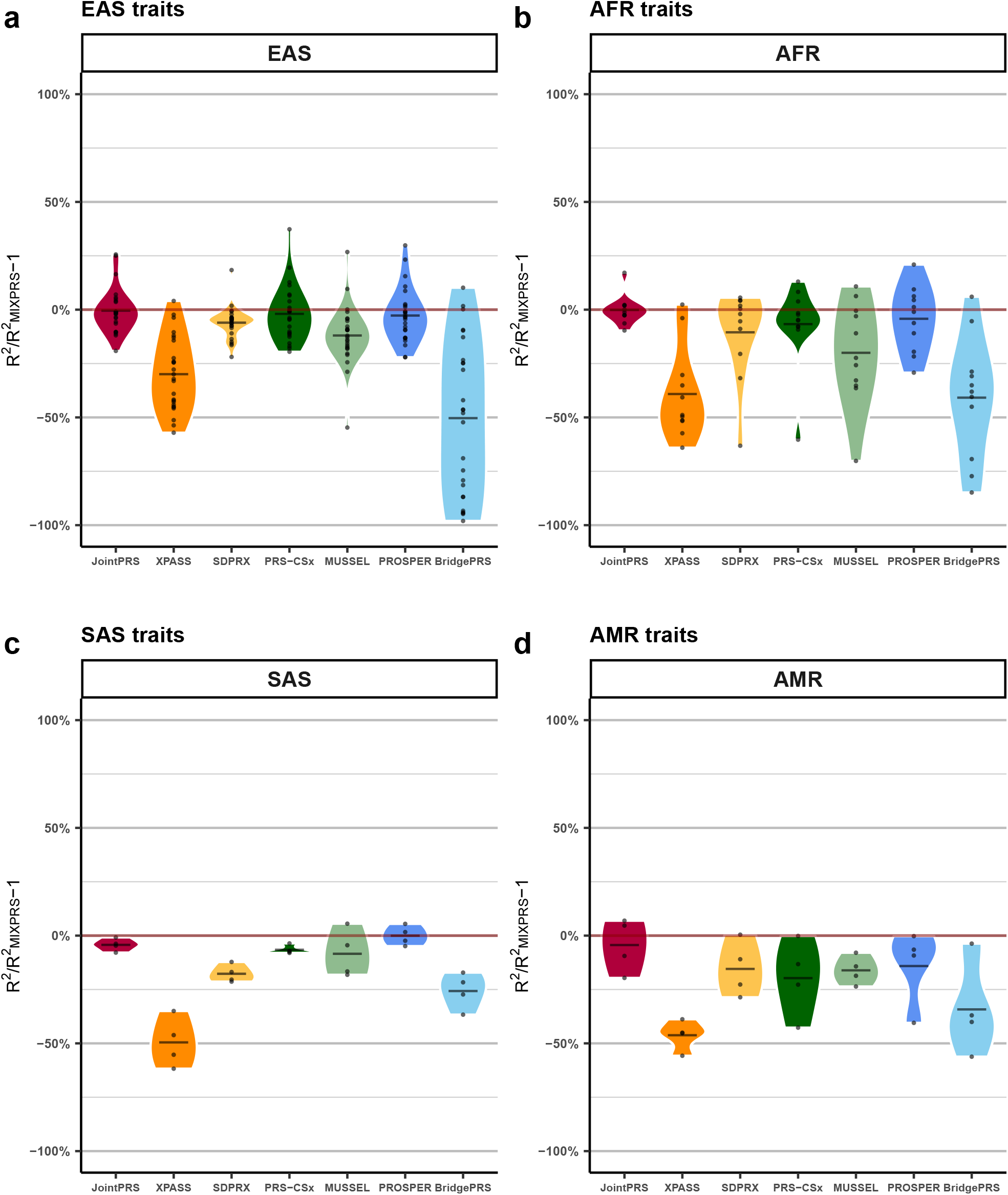
Relative prediction accuracy of multi-population PRS methods compared to MIXPRS across 26 traits when tuning and testing data are from the same cohort (UKBB). **a–d**, Relative prediction accuracy was evaluated across 26 traits in four non-European populations (**a** EAS; **b** AFR; **c** SAS; and **d** AMR), using tuning and testing data from UKBB. Performance was assessed through a 5-fold cross-validation within UKBB. Relative performance of seven existing methods compared to MIXPRS was measured as 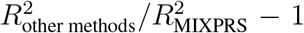 for quantitative traits and AUC_other methods_*/*AUC_MIXPRS_ − 1 for binary traits across the five folds. Results are presented as violin plots, with the mean across traits indicated by a black crossbar within each violin.

**Figure 6.**
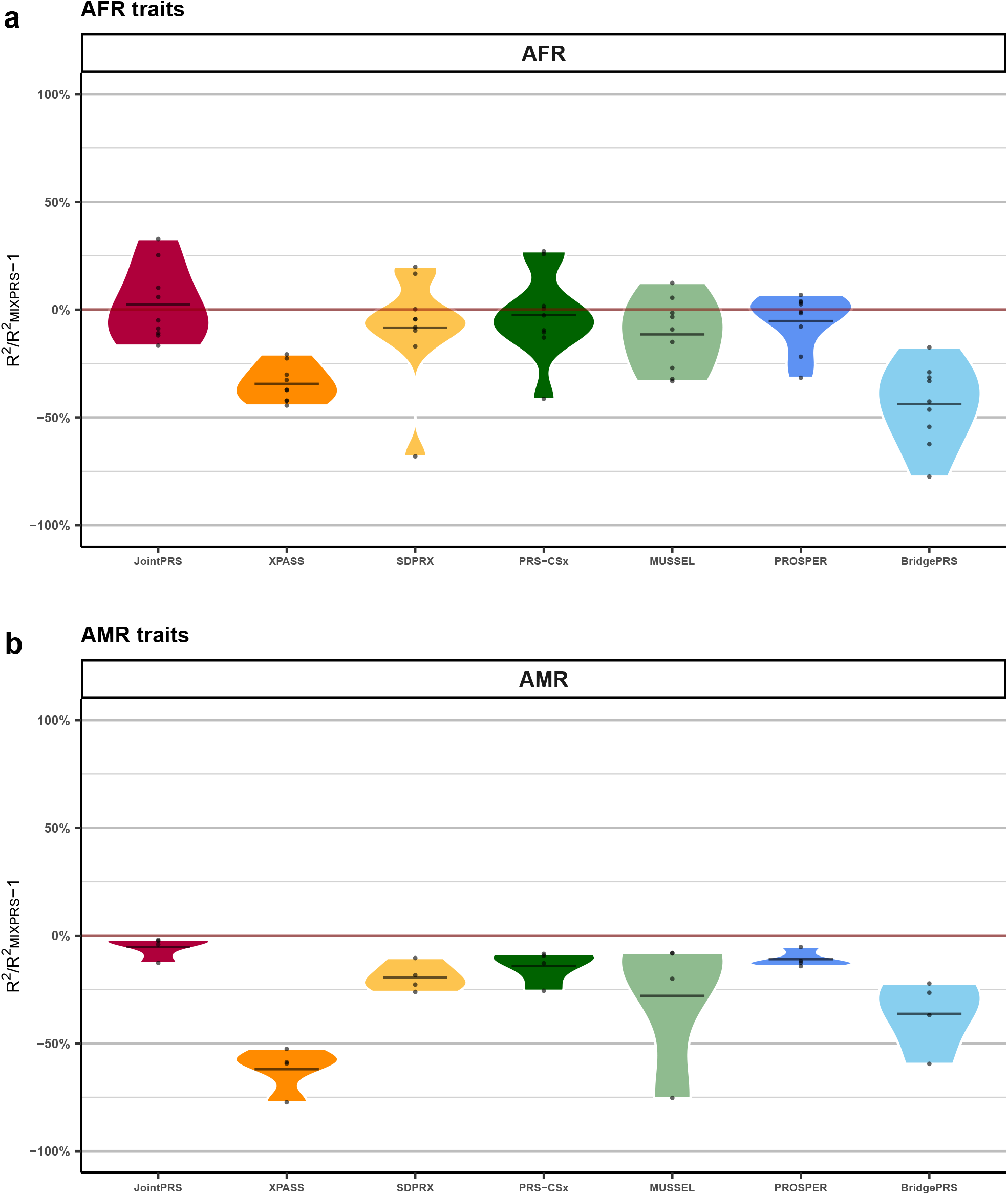
Relative prediction accuracy of multi-population PRS methods compared to MIXPRS across nine traits when tuning and testing data are from different cohorts (UKBB and AoU). **a–b**, Relative prediction accuracy was assessed across nine traits in two non-European populations (**a** AFR; **b** AMR) using tuning data from UKBB and testing data from AoU. The relative performance of seven existing methods compared to MIXPRS was calculated as 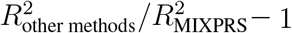 for quantitative traits. Results are presented as violin plots, with the mean relative performance across traits indicated by a black crossbar within each violin.

Additionally, we evaluated the predictive performance of MIXPRS, IndPRS, and the component method SDPRX in the AoU dataset. IndPRS integrates PRS across methods and populations by leveraging individual-level UKBB data for tuning, followed by evaluation in AoU. As shown in Figure S9 and Table S18, MIXPRS achieved predictive performance comparable to IndPRS, suggesting minimal information loss when combining PRS using only GWAS summary statistics instead of individual-level data. Both MIXPRS and IndPRS significantly outperformed SDPRX, underscoring the advantage of integrating PRS across multiple methods and populations. In conclusion, MIXPRS eliminates the necessity for individual-level tuning data while achieving top-tier performance across various data scenarios.

## Discussion

In this study, we introduced MIXPRS, an efficient multi-population PRS integration framework designed to combine PRS from multiple methods and diverse populations using only GWAS summary statistics. By employing strategies such as SNP pruning to mitigate the LD mismatch issue and NNLS to ensure robust estimation of PRS combination weights, MIXPRS significantly enhances the accuracy and robustness of PRS predictions. We benchmarked MIXPRS against seven existing multi-population methods: JointPRS, XPASS, SDPRX, PRS-CSx, MUSSEL, PROSPER, and BridgePRS as well as one single-population PRS integration framework: PUMAS-EN. Through extensive simulations and real-data analyses across 22 continuous and four binary traits in the UKBB and AoU datasets, we demonstrated that MIXPRS consistently achieves superior predictive accuracy.

Traditional PRS methods often rely heavily on individual-level data for model tuning, hyper-parameter selection, and PRS integration, introducing potential biases, instability, and reduced generalizability due to small tuning sample sizes [19, 20, 21, 22]. Our approach, complemented by recent developments in PRS tuning and integration frameworks, effectively addresses these limitations by adapting PRS methods to exclusively utilize large GWAS summary statistics [31, 32, 33, 47]. This adaptation mitigates privacy concerns associated with individual-level genetic data, eliminates biases and noise introduced by small tuning datasets, and facilitates equitable, standardized benchmarking by relying exclusively on publicly available GWAS summary statistics.

One critical issue addressed in this work is the LD mismatch, which significantly impacts genetic analyses but has received limited attention in the existing literature. Only a few prior studies have proposed methods to mitigate this issue [48]. We explicitly quantified the impact of LD mismatch within the PRS integration framework by analyzing residual correlations from pseudo-GWAS subsampling and the final PRS prediction accuracy. Our results revealed that LD mismatch substantially contributes to high residual correlations, leading to overfitting and diminished predictive accuracy. While we proposed SNP pruning as a practical mitigation strategy, alternative approaches include shrinkage-based regularization of the LD matrix, such as the Ledoit–Wolf estimator [49], and SNP filtering based on conditional distribution, as implemented in DENTIST [50]. Further research is needed to systematically assess the broader impact of LD mismatch and to develop robust, widely applicable solutions across diverse genetic study designs.

We note two limitations of MIXPRS has two primary limitations that need further investigation. First, our current implementation integrates only two multi-population methods, JointPRS-auto and SDPRX, which do not require individual-level tuning data. Integrating additional multi-population PRS methods that rely on individual-level data may be further explored [31, 32, 33, 47]. Prior studies suggest individual-level tuning processes could be adapted for use within a GWAS summary statistics-only framework, necessitating additional research to effectively incorporate these multi-population methods within the current MIXPRS framework [31, 32, 33, 47]. Second, although we utilized NNLS for estimating PRS combination weights, alternative regression methods such as ridge, lasso, and elastic net, may also be considered to see whether they may lead to better performance [33].

The PRS integration framework presented here is not restricted to combining methods and populations. Future research should explore the potential benefits of integrating PRS derived from additional information sources, such as PRS for related traits, to further enhance predictive accuracy [51, 52, 53, 54].

## Methods

### MIXPRS method

#### Step1: GWAS subsampling in MIXPRS

This section describes the GWAS subsampling method in MIXPRS, specifically addressing: (i) derivation of GWAS summary statistics, (ii) data fission and the ideal pseudo-GWAS subsampling procedure, (iii) overfitting issues arising from LD mismatch in practical pseudo-GWAS subsampling, and (iv) LD pruning strategy implemented in MIXPRS to effectively mitigate LD mismatch.

**(i) Derivation of GWAS summary statistics**. Based on the additive genetic model for the target population *k*, we have:

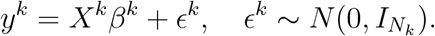

Here, *y*^*k*^, *X*^*k*^, *β*^*k*^, *ϵ*^*k*^, and *N*_*k*_ represent the standardized phenotype vector, the column-standardized genotype matrix, the standardized SNP effect-size vector, the residual vector, and the sample size for population *k*, respectively. The GWAS summary statistics 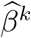 for population *k* are obtained as marginal least-squares estimates of SNP effects:

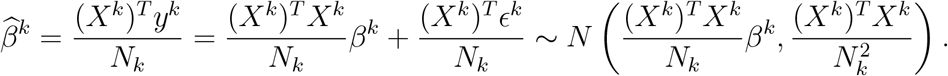

We denote the LD pattern and new residuals for population *k* by:

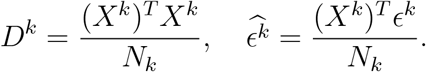

Thus, the GWAS summary statistics can be concisely expressed as:

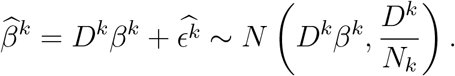

**(ii) Data fission and the ideal pseudo-GWAS subsampling procedure**. To generate two independent GWAS summary statistics, 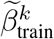 and 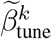, from the original GWAS summary statistics 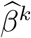, we utilize the following data fission theorem for Gaussian variables [28]:

**Theorem 1** (Data Fission for Gaussian Variables). *Suppose X ∼ N* (*µ*, Σ) *is a d-dimensional Gaussian vector* (*d ≥* 1). *Independently draw Z ∼ N* (0, Σ), *and let τ ∈* (0, *∞*) *be a tuning parameter. Define:*

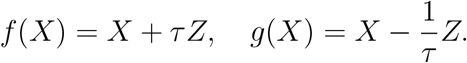

*Then, the following properties hold:*

- *f(X) ∼ N (µ, (*1 *+ τ*^2^)Σ).
- *g*(*X*) *∼ N* (*µ*, (1 + *τ*^−2^)Σ).
- *f* (*X*) *and g*(*X*) *are independent random variables*.
- *A larger value of τ implies that f* (*X*) *is less informative, and g*(*X*)|*f* (*X*) *is correspondingly more informative*.

Now, applying Theorem 1 with 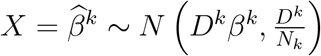, independently draw 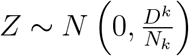, we generate two independent GWAS summary statistics:

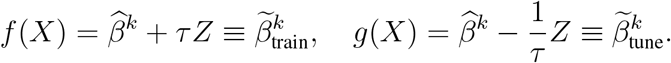

Therefore,

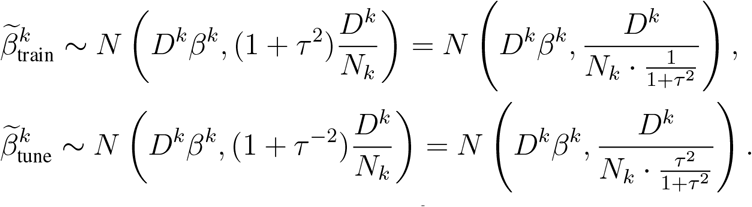

Define 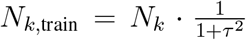 and 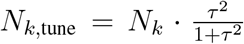, noting that *N*_*k*,train_ + *N*_*k*,tune_ = *N*_*k*_. Let 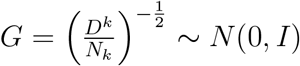, the above expressions can be equivalently written as:

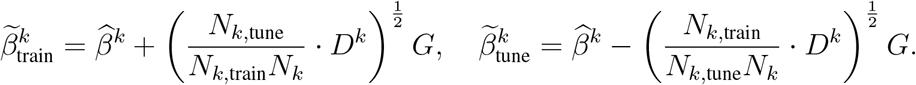

This is the ideal pseudo-GWAS subsampling procedure.

Note that in practice, following PUMAS-EN [33], we set the ratio of subsampled training to tuning GWAS sample sizes as *N*_*k*,train_*/N*_*k*,tune_ = 3, and apply a 4-fold Monte Carlo cross-validation (MCCV), repeating the GWAS subsampling procedure four times.

**(iii) Overfitting issues arising from LD mismatch in practical pseudo-GWAS subsampling**. In practice, the true LD pattern *D*^*k*^ is unknown, and we instead use an external LD reference panel 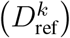, such as from the 1000 Genomes Project, to perform pseudo-GWAS subsampling:

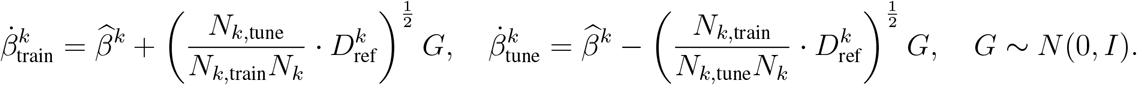

For the ideal pseudo-GWAS subsampling, we have:

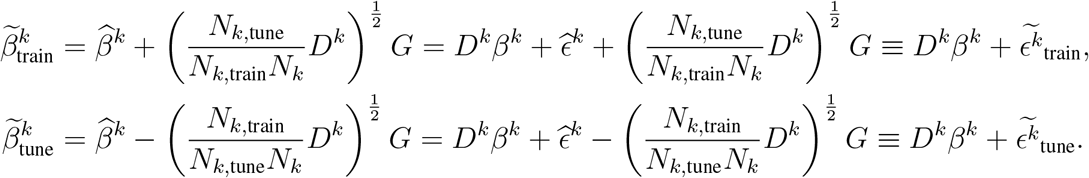

Thus, the residuals follow:

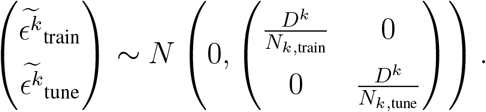

In contrast, for the practical pseudo-GWAS subsampling procedure:

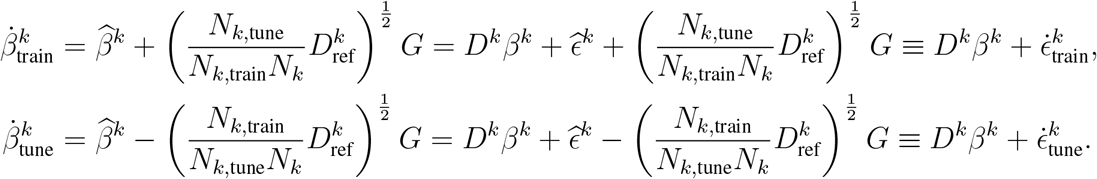

The residuals now follow a correlated structure:

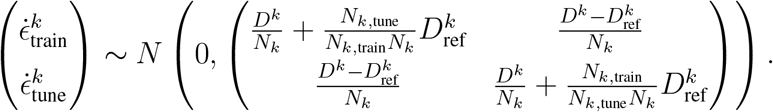

Consequently, the ideal pseudo-GWAS subsampling produces residuals with zero correlation between training and tuning residuals, whereas the practical pseudo-GWAS subsampling yields non-zero residual correlations due to LD mismatch. This residual correlation potentially leads to overfitting in subsequent PRS integration steps.

**(iv) LD pruning strategy implemented in MIXPRS to effectively mitigate LD mismatch**. To address potential overfitting induced by LD mismatch, MIXPRS implements an LD pruning strategy. LD mismatch occurs due to slight variations in LD structures across genetic cohorts, which can accumulate significantly given the involvement of millions of SNPs, leading to substantial residual correlations and potential overfitting. Although the exact LD patterns differ slightly between cohorts, the overall LD structure within the same population generally remains consistent. Leveraging this consistency, MIXPRS selects approximately independent SNPs by retaining variants that exhibit pairwise correlation below 0.5 within sliding windows of 250 kb. These LD-pruned SNPs are identified using PLINK [36] with a reference genotype panel matched to the target population, such as the 1000 Genomes Project. Subsequently, MIXPRS restricts the GWAS subsampling procedure exclusively to these LD-pruned SNPs and replaces the unknown LD matrix *D*^*k*^ with an identity covariance matrix (*I*). This approach significantly alleviates LD mismatch, enhancing robustness and mitigating overfitting in the GWAS subsampling and subsequent PRS integration steps.

#### Step2: Estimation of MIXPRS combination weights for different PRS methods

This section outlines the estimation of the weights for different PRS methods in MIXPRS, which consists of two substeps: (i) generating LD-pruned PRS across methods and populations, and (ii) estimating the combining weights used to integrate these LD-pruned PRS.

A. **Generating LD-pruned PRS across methods and populations** Starting from the LD-pruned, subsampled training GWAS summary statistics 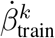 for the target population *k*, we first harmonize each other population’s GWAS summary statistics 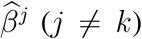 to the same pruned SNP set. These aligned, pruned GWAS summary statistics across all populations then feed into JointPRS-auto and SDPRX, yielding per-population LD-pruned PRS effect-size vectors: 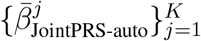 and 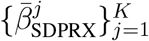. Note that JointPRS-auto produces all *K* populations’ PRS in one joint model. SDPRX, by contrast, pairs each non-European GWAS in turn with the European summary statistics to generate non-European PRS (and uses the paired target and European GWAS summary statistics for the European output). To stabilize our final combining weights, we repeat this entire PRS-derivation step four times using each of the four pseudo-GWAS training subsamples, then average the results.
B. **Estimating the combining weights used to integrate these LD-pruned PRS**. We first estimate non-negative combining weights from individual-level tuning data 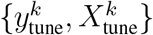. Define the PRS feature matrix

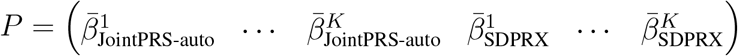

We then solve

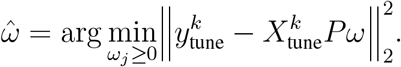

via the active-set Lawson–Hanson NNLS algorithm [37]. To eliminate the need for individual-level tuning data, we reformulate the Lawson–Hanson NNLS algorithm entirely in summary-statistic form, leveraging the LD-pruned, subsampled tuning GWAS 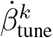 and the LD matrix *D*^*k*^ (see Algorithm 1). We apply this summary-statistic NNLS procedure independently to each of the four pseudo-GWAS tuning subsamples.

##### Algorithm 1

Lawson–Hanson NNLS for Estimating Non-Negative PRS Combining Weights

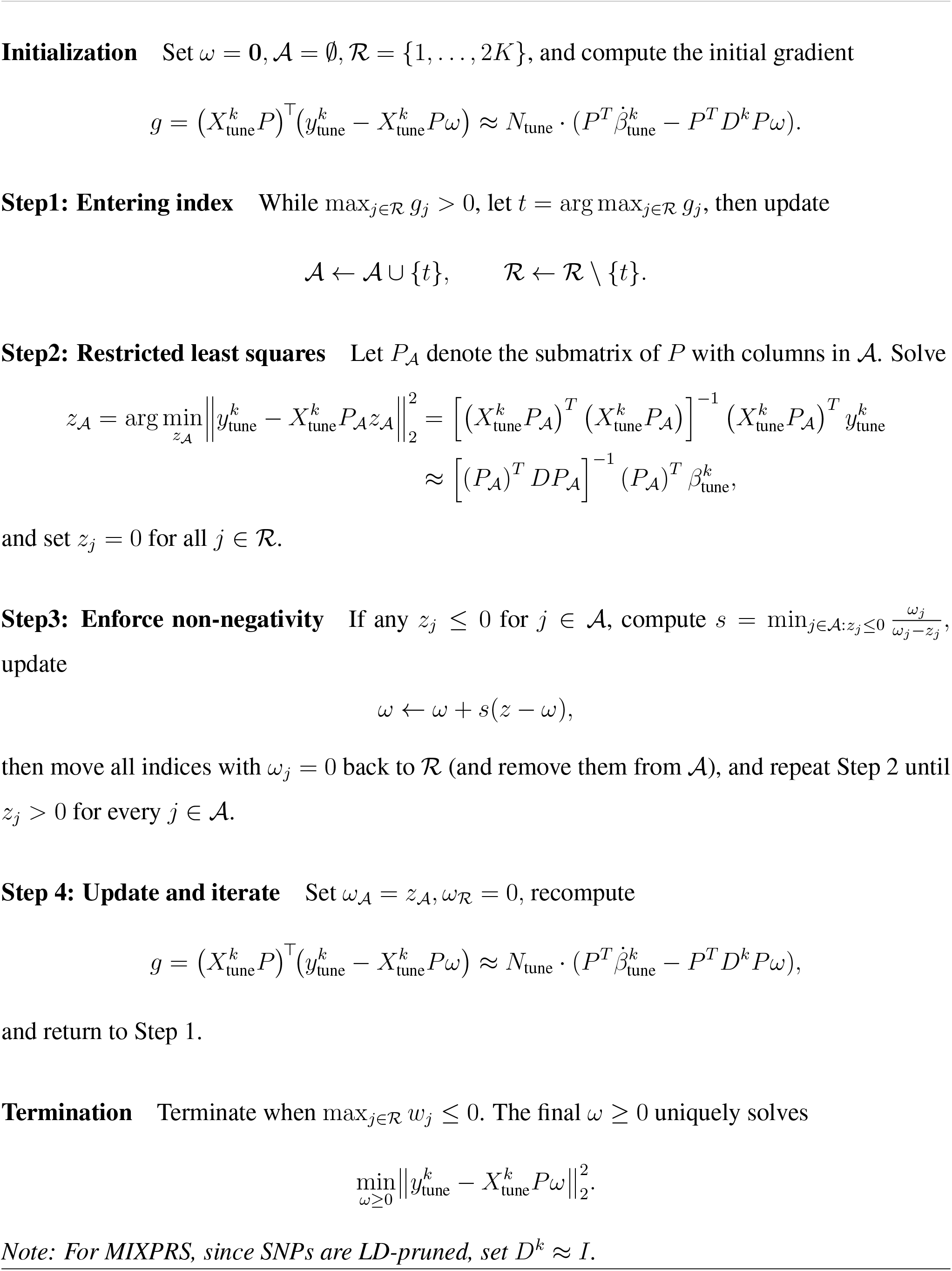

#### Step3: Derivation of MIXPRS

Building on the non-negative weight estimates 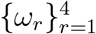 obtained from step2 (one for each pseudo-GWAS subsample), we define the final PRS combining weight vector as:

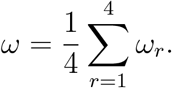

We then apply JointPRS-auto and SDPRX to the original GWAS summary statistics from all K populations, yielding the PRS effect-sizes estimates:

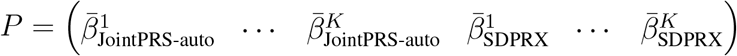

The final MIXPRS for the target population k is obtained as:

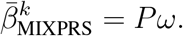

### Existing PRS methods

#### JointPRS(−auto)

JointPRS [14] integrates GWAS summary statistics and LD reference panels from multiple populations through a shared Bayesian shrinkage model incorporating genetic correlation structures. When individual-level tuning data are unavailable, JointPRS-auto directly estimates genetic correlations and shrinkage parameters using GWAS summary statistics alone. When individual-level tuning data are available, JointPRS computes two versions: meta version, which automatically estimates shrinkage parameters; tune version, which optimizes shrinkage parameters and performs linear combination across populations. The optimal version (meta or tune) is then selected using a data-adaptive strategy based on predictive accuracy evaluated in the individual-level tuning dataset.

#### XPASS

XPASS [15] jointly integrates GWAS summary statistics and LD structures from two populations using a bivariate Gaussian distribution to model genetic correlation, facilitating information transfer from an auxiliary population to a target population. It also incorporates population-specific effects identified through a P+T procedure. Importantly, XPASS does not require individual-level tuning data.

#### SDPRX

SDPRX [18] employs a hierarchical Bayesian framework to model GWAS summary statistics and LD structures from two populations. It categorizes SNP effects into none, population-specific, or shared across populations. The shared component is modeled via mixtures of bivariate Gaussian distributions informed by genetic correlation. SDPRX does not require individual-level tuning data.

#### PRS-CSx(−auto)

PRS-CSx [17], an extension of PRS-CS [41], integrates multi-population GWAS summary statistics via a shared Bayesian shrinkage prior. In the absence of tuning data, PRS-CSx automatically estimates a global shrinkage parameter through a Bayesian approach. When tuning data are available, it selects the best-performing shrinkage parameter from a prede-fined set and performs linear combination across populations, optimizing prediction accuracy.

#### MUSSEL

MUSSEL [20] integrates GWAS summary statistics and LD structures across multiple populations using a multivariate spike-and-slab prior capturing genetic correlation. This method necessitates individual-level tuning data to determine parameters including causal SNP proportion, heritability within populations, and between-population correlations. A subsequent super-learning step selects from linear methods (lasso, ridge, elastic net, and linear regression) to further optimize predictions.

#### PROSPER

PROSPER [21] utilizes a penalized linear regression framework, combining Lasso and ridge penalties to account for genetic sparsity and population similarities. Individual-level tuning data are required to select optimal penalty parameters. An additional super-learning step selects from linear methods (lasso, ridge, elastic net, and linear regression) to integrate PRS generated across penalty parameters and populations, further enhancing predictive performance.

#### BridgePRS

BridgePRS [22] incorporates GWAS summary statistics from two populations by modeling shared and population-specific SNP effects. Initially, it applies Gaussian priors to estimate SNP effects separately for each population. Subsequently, it integrates auxiliary population information to refine effect-size estimates in the target population. Individual-level tuning data are required to optimally combine these estimates into a final PRS score using ridge regression.

#### PUMAS-EN

PUMAS-EN [33] is a single-population PRS integration framework designed to integrate PRS across multiple populations and methods through an ensemble learning approach using GWAS summary statistics and LD reference panels from various populations.

Two versions were considered in our evaluation: the first version, PUMAS-EN, utilized only the 1000 Genomes Project LD reference panel throughout all integration stages, including GWAS subsampling, PRS training, and ensemble learning, closely matching the MIXPRS setup. This version incorporated four commonly used single-population PRS methods: lassosum [39], LD-pred2 [40], PRS-CS [41], and SBLUP [42]. The second version, PUMAS-EN paper, used the 1000 Genomes Project LD reference panel only for GWAS subsampling but employed UKBB genotype data for PRS training and ensemble learning. This version included a broader set of nine single-population PRS methods: lassosum, LDpred2, PRS-CS, MegaPRS [43], SBayesR [44], DBSLMM [45], Vilma [46], and SBLUP.

Both methods subsequently integrate these diverse PRS models via a penalized regression approach optimized using 4-fold MCCV. Ensemble parameters are tuned with subsampled GWAS summary statistics, thus removing any requirement for individual-level data. The final ensemble model weights are determined by averaging coefficients across MCCV folds.

### Simulation design

#### Simulation setting

We simulated the standardized true effect sizes using a spike-and-slab model for five populations: EUR, EAS, AFR, SAS, and AMR:

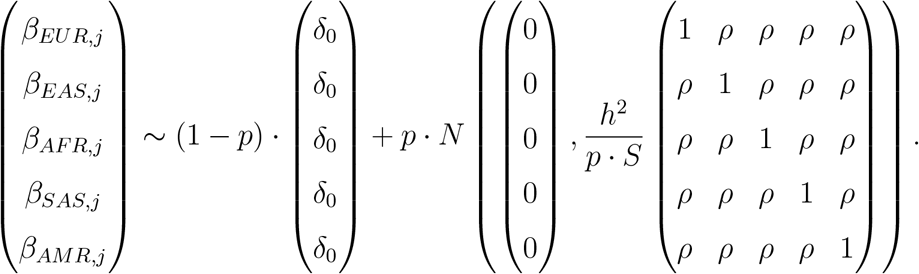

Simulations were performed under a genetic architecture with heritability fixed at *h*^2^ = 0.4, varying causal SNP proportions (*p* = 0.1, 0.01, 0.001, 5 *×* 10^−4^), and cross-population genetic correlations of *ρ* = 0.8, across 1,203,063 HapMap3 SNPs. Genotype data for the European population were obtained from the UKBB (*N* = 311, 600) [34], while non-European genotype data were obtained from publicly available simulated genotype datasets based on 1000 Genomes Project (*N* = 25, 000 and 90, 000) [19]. For each simulation setting, we generated five replicates of the effect sizes.

After simulating the true effect sizes across difference scenarios, we used GCTA-sim [55] to generate standardized phenotypes *y* based on the heritability *h*^2^, the simulated standardized true effect sizes *β*, and column-standardized genotype datasets *X*, under an addictive genetic model:

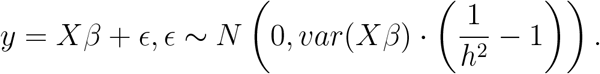

We then employed PLINK2 [56] to derive GWAS summary statistics from simulated phenotypes and genotype datasets. To ensure fair comparisons across methods, we extracted the set of 717,985 SNPs that were common across the LD reference panels of all methods and populations. This intersection ensures that all methods used the same set of SNPs in GWAS.

For methods requiring individual-level tuning data (JointPRS, PRS-CSx, MUSSEL, PROSPER, and BridgePRS), training GWAS sample sizes were set at 15,000 and 80,000, respectively, with 10,000 individuals reserved for parameter tuning. For methods utilizing only GWAS summary statistics (XPASS and SDPRX), training GWAS sample sizes were set to 25,000 and 90,000. Each simulation scenario was replicated five times, with detailed simulation procedures provided in the Methods section.

#### Simulation residual correlation analysis

As detailed in the Methods section (MIXPRS method, Step1: GWAS subsampling in MIXPRS, (iii): Overfitting issues arising from LD mismatch in practical pseudo-GWAS subsampling), we observed correlation between training and tuning residuals resulting from the practical pseudo-GWAS subsampling procedure:

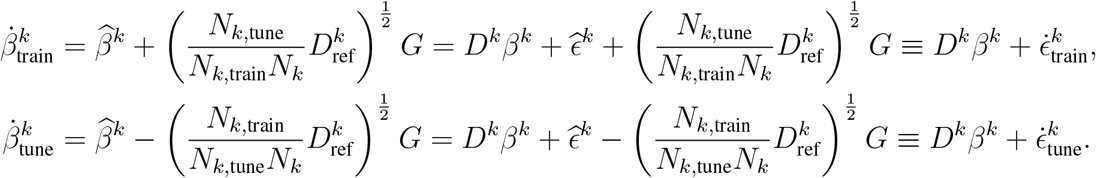

The residuals now follow a correlated structure:

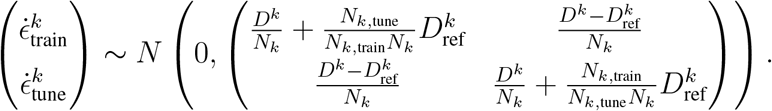

We denote 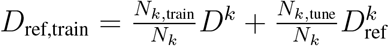, and 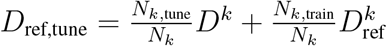, then we have

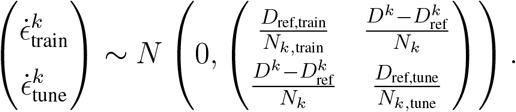

In simulations, as we know the true effect sizes *β*^*k*^ and the LD pattern of GWAS summary statistics *D*^*k*^, we calculate residuals 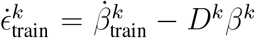 and 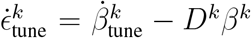. We further calculate the standardized residuals 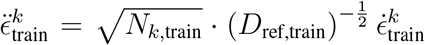, and 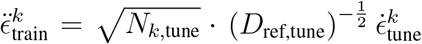. The residual correlation is then defined as the Pearson correlation coefficient between the standardized training and tuning residual vectors across all *J* SNPs, calculated as:

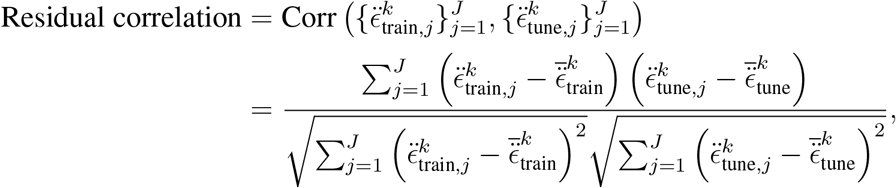

where 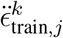 and 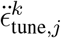 denote the standardized residuals for SNP *j* from the training and tuning datasets, respectively, and 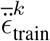 and 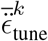 represent their corresponding averages across all SNPs.

### Real-data analysis

We confirm that this research complies with all relevant ethical regulations. Participants from the UK Biobank provided written informed consent (further details available at https://www.ukbiobank.ac.uk/learn-more-about-uk-biobank/governance). Data from participants in the All of Us Research Program were collected according to the All of Us Research Program Operational Protocol (https://allofus.nih.gov/article/all-us-research-program-protocol), with consent procedures detailed at https://allofus.nih.gov/about/protocol/all-us-consent-process.

### GWAS summary statistics

We compiled GWAS summary statistics for 22 continuous and four binary traits across five populations (EUR, EAS, AFR, SAS, and AMR) from multiple consortia [5, 6, 7, 8, 9, 11, 13, 57, 58, 59, 60, 61, 62, 63, 64, 65, 66]. Quality control followed LDHub guidelines using LDSC to exclude duplicate SNPs, strand-ambiguous SNPs (A/T and G/C), insertions and deletions (INDELs), and SNPs with effective sample sizes less than 0.67 times the 90th percentile threshold [67, 68]. To facilitate fair comparisons across methods while accommodating population-specific genetic variants, SNP sets for each population were further restricted to variants consistently available across all evaluated methods within the corresponding population. The 1000 Genomes Project served as the LD reference panel. Comprehensive information on GWAS summary statistics is provided in Table S1 and Table S2. Given our primary objective to evaluate prediction accuracy specifically in non-European populations (EAS, AFR, SAS, and AMR), we verified that no overlap exists between training GWAS datasets and individual-level tuning or testing datasets derived from the UKBB or AoU cohorts in these populations. Although overlaps may exist between EUR training GWAS and corresponding tuning or testing datasets, any resulting bias is considered negligible for the evaluation within non-European populations.

### UKBB data

For the UKBB dataset [34], we classified individuals into five super-populations using the ancestry inference procedure implemented in SDPRX [18]. Specifically, we performed principal-component analysis (PCA) jointly on UKBB participants and 1000 Genomes Project reference samples and subsequently trained a random forest classifier based on the top ten principal components to assign ancestry labels. The resulting population counts were EUR (311,600), EAS (2,091), AFR (6,829), SAS (7,857), and AMR (635).

We obtained data for 22 quantitative phenotypes from UKBB participants; detailed descriptions are provided in Table S3. For systolic blood pressure (SBP) and diastolic blood pressure (DBP), manual and automated measurements were combined following established guidelines [59]. Participants lacking medication records (fields 6177 and 6153) were excluded from analyses of low-density lipoprotein (LDL), total cholesterol (TC), SBP, and DBP, since medication status is necessary for accurate adjustment of these traits, consistent with previous studies [5, 59].

Case-control status for four binary traits was determined using ICD-9 and ICD-10 diagnostic codes, operation codes, and self-reported disease information. For breast cancer, analyses included only female participants. Effective sample sizes for binary traits were computed as:

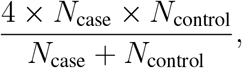

where *N*_case_ and *N*_control_ denote the number of cases and controls, respectively.

### AoU data

For the AoU dataset [35], participants were classified into six populations (EUR, EAS, AFR, SAS, AMR, and Middle Eastern (MID)) using the procedure and reference samples described by Venner et al. [69]. Population-specific counts were EUR (133,581), EAS (5,706), AFR (56,913), SAS (3,217), AMR (45,035), and MID (942). Our analyses specifically focus on prediction accuracy within AFR and AMR populations.

We extracted nine quantitative phenotypes from AoU participants using the concept IDs detailed in Table S4. Height and body mass index (BMI) values were directly utilized, whereas for the remaining seven traits, we calculated the median of multiple measurements per individual. For all nine traits, we excluded outliers defined as observations satisfying:

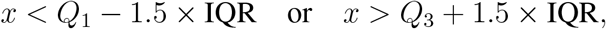

where *Q*_1_ and *Q*_3_ represent the first and third quartiles, respectively, and IQR = *Q*_3_ − *Q*_1_.

### Evaluation metrics

For 22 quantitative traits, predictive accuracy was assessed using *R*^2^:

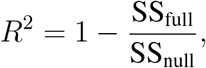

where SS_null_ and SS_full_ denote the residual sums of squares from the null model (including age, sex, and the first 20 genetic principal components) and the full model (additionally including the PRS), respectively. For four binary traits, predictive accuracy was assessed using logistic regression, quantified by the area under the receiver operating characteristic curve (AUC).

To facilitate comparisons between methods, we defined the relative improvement of method B over method A as:

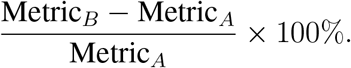

## Data availability

The GWAS summary statistics datasets used in this study are publicly accessible as detailed below. The BBJ GWAS summary-level statistics data are available in the NBDC Human Database under accession codes hum0014-v36 [https://humandbs.dbcls.jp/en/hum0014-v36] and hum0197-v23 [https://humandbs.dbcls.jp/en/hum0197-v23]. The BCAC GWAS summary-level statistics data are available in the GWAS Catalog under accession code 29059683 [https://www.ebi.ac.uk/gwas/publications/29059683]. The BCX GWAS summary-level statistics data are available in the GWAS Catalog under accession code 32888493 [https://www.ebi.ac.uk/gwas/publications/32888493]. The CARDIoGRAM GWAS summary-level statistics data are available in the GWAS Catalog under accession code 21378990 [https://www.ebi.ac.uk/gwas/publications/21378990]. The TRICL-ILCCO and LC3 GWAS summary-level statistics data are available in the GWAS Catalog under accession code 28604730 [https://www.ebi.ac.uk/gwas/publications/28604730]. The DIAGRAM GWAS summary-level statistics data are available in the GWAS Catalog under accession code 28566273 [https://www.ebi.ac.uk/gwas/publications/28566273]. The GIANT GWAS summary-level statistics data are available from the GIANT consortium [https://portals.broadinstitute.org/collaboration/giant/index.php/GIANT_consortium_data_files]. The GLGC GWAS summary-level statistics data are available from the Global Lipids Genetics Consortium [https://csg.sph.umich.edu/willer/public/glgc-lipids2021/]. The ICBP GWAS summary-level statistics data are available in the GWAS Catalog under accession code 30224653 [https://www.ebi.ac.uk/gwas/publications/30224653]. The PAGE GWAS summary-level statistics data are available in the GWAS Catalog under accession code 31217584 [https://www.ebi.ac.uk/gwas/publications/31217584]. The UKBB Liver Enzymes GWAS summary-level statistics data are available in the GWAS Catalog under accession code 33972514 [https://www.ebi.ac.uk/gwas/publications/33972514].

Access to individual-level data from the UK Biobank (UKBB) can be requested through the UKBB data access portal [https://www.ukbiobank.ac.uk/enable-your-research/apply-for-access]. Access to individual-level data from the All of Us (AoU) Research Program can be requested through the All of Us data access portal [https://www.researchallofus.org/data-tools/data-access/].

## Code availability

The code for all simulation studies and real-data analyses presented in this paper is publicly available on GitHub at https://github.com/LeqiXu/MIXPRS_analysis. Software implementing the proposed MIXPRS method is available at https://github.com/YCSGP/MIXPRS.

For comparative analyses, implementations of other methods are accessible via their respective repositories: JointPRS at https://github.com/YCSGP/JointPRS, XPASS at https://github.com/YangLabHKUST/XPASS, SDPRX at https://github.com/eldronzhou/SDPRX, PRS-CSx at https://github.com/getian107/PRScsx, MUSSEL at https://github.com/Jin93/MUSSEL, PROSPER at https://github.com/Jingning-Zhang/PROSPER, BridgePRS at https://github.com/clivehoggart/BridgePRS, PUMAS-EN at https://github.com/qlu-lab/PUMAS.

## Acknowledgments

This research was supported in part by NIH grant R01 HG012735 and NSF grant DMS 2310836. We thank Jiaqi Hu, Chen Lin, Dr. Lijun Wang, and Dr. Yingxin Lin for their invaluable discussions. We are also grateful to Dr. Qiongshi Lu and Stephen Dorn for sharing their codes, LD reference panels, and for their suggestions on implementing various methods.

Our research utilized the UK Biobank resource under approved data request (refs: 29900) and the All of Us resource. We gratefully acknowledge UK Biobank and All of Us participants for their contributions, without whom this research would not have been possible. We also thank the National Institutes of Health’s All of Us Research Program for making available the participant data examined in this study.

## Author contributions

H.Z. and L.X. conceived and designed the study. L.G. and L.X. developed the statistical framework and algorithm. L.X. conducted all simulations and real-data analyses under the supervision of H.Z. X.Z. assisted with simulations. Y.D. assisted with exploratory analyses. Y.D., G.Z., and Z.B. assisted with real-data analyses. L.X. developed the MIXPRS software. L.X., Y.D., X.Z., Z.B., G.Z., L.G., and H.Z. drafted or substantively revised the manuscript. All authors reviewed and approved the final version of the manuscript.

