## Supplemental Information for "Almost Free Enhancement of Multi-Population PRS: From Data-Fission to Pseudo-GWAS Subsampling"

Table S14: Real data results of MIXPRS, PUMAS-EN, and PUMAS-EN_paper for $R^2$ of four lipids traits in EAS, AFR, and SAS UKBB without tuning data (UKBB) . 26

### 1 Supplemental Figures

**Figure S1. Prediction accuracy of MIXPRS and its component methods (JointPRS-auto and SDPRX) across varying genetic architectures in simulations. a–b**, Simulations were conducted with total heritability fixed at $h^2 = 0.4$, cross-population genetic correlation fixed at $\rho = 0.8$, and four causal SNP proportions ($p = 0.1, 0.01, 0.001, 5 \times 10^{-4}$). Non-European populations (EAS, AFR, SAS, and AMR) had a fixed training sample size of $N_{\text{train}} = 100,000$, whereas the EUR training sample size was fixed at $N_{\text{train}} = 311,600$. Prediction accuracy was evaluated using $R^2$. Results are shown as **a** dot plots and **b** bar plots, where each dot or bar represents the mean prediction accuracy across five simulation replicates for each scenario. The best-performing and second-best-performing methods are indicated by two stars and one star, respectively, above the corresponding bars.

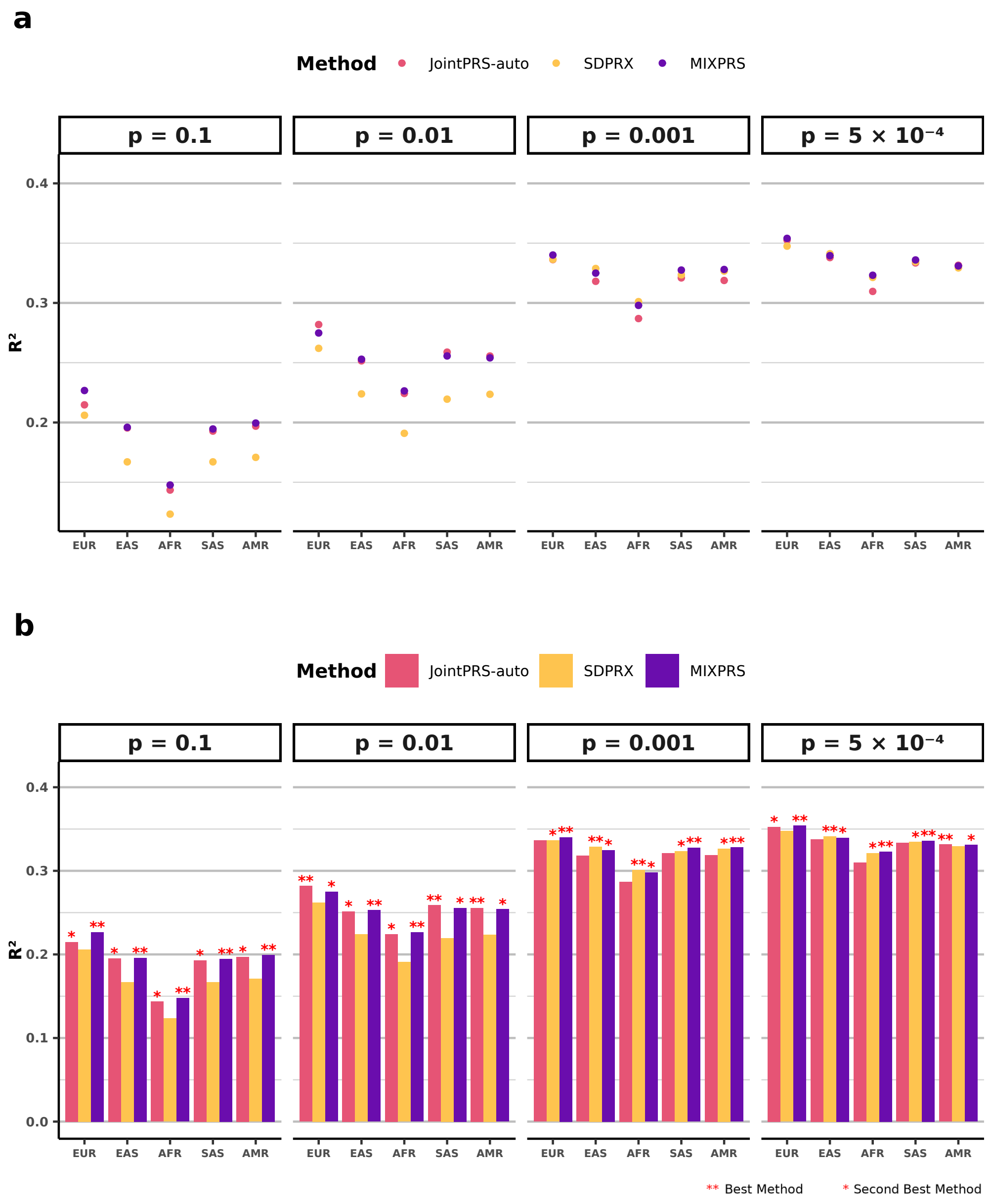

**Figure S2. Residual correlation and prediction accuracy of MIXPRS under different GWAS subsampling strategies across genetic architectures in simulations. a–b**, Simulations were performed with total heritability fixed at $h^2 = 0.4$, cross-population genetic correlation fixed at $\rho = 0.8$, and four causal SNP proportions ($p = 0.1, 0.01, 0.001, 5 \times 10^{-4}$). Non-European populations (EAS, AFR, SAS, and AMR) had a fixed training sample size of $N_{\text{train}} = 100,000$, whereas the EUR training sample size was fixed at $N_{\text{train}} = 311,600$. **a** Residual correlations between subsampled training and tuning GWAS datasets for three GWAS subsampling strategies, with each dot representing the mean residual correlation across five simulation replicates per scenario. **b** Prediction accuracy ($R^2$) of MIXPRS across the three GWAS subsampling strategies, with each dot representing the mean prediction accuracy across five simulation replicates per scenario.

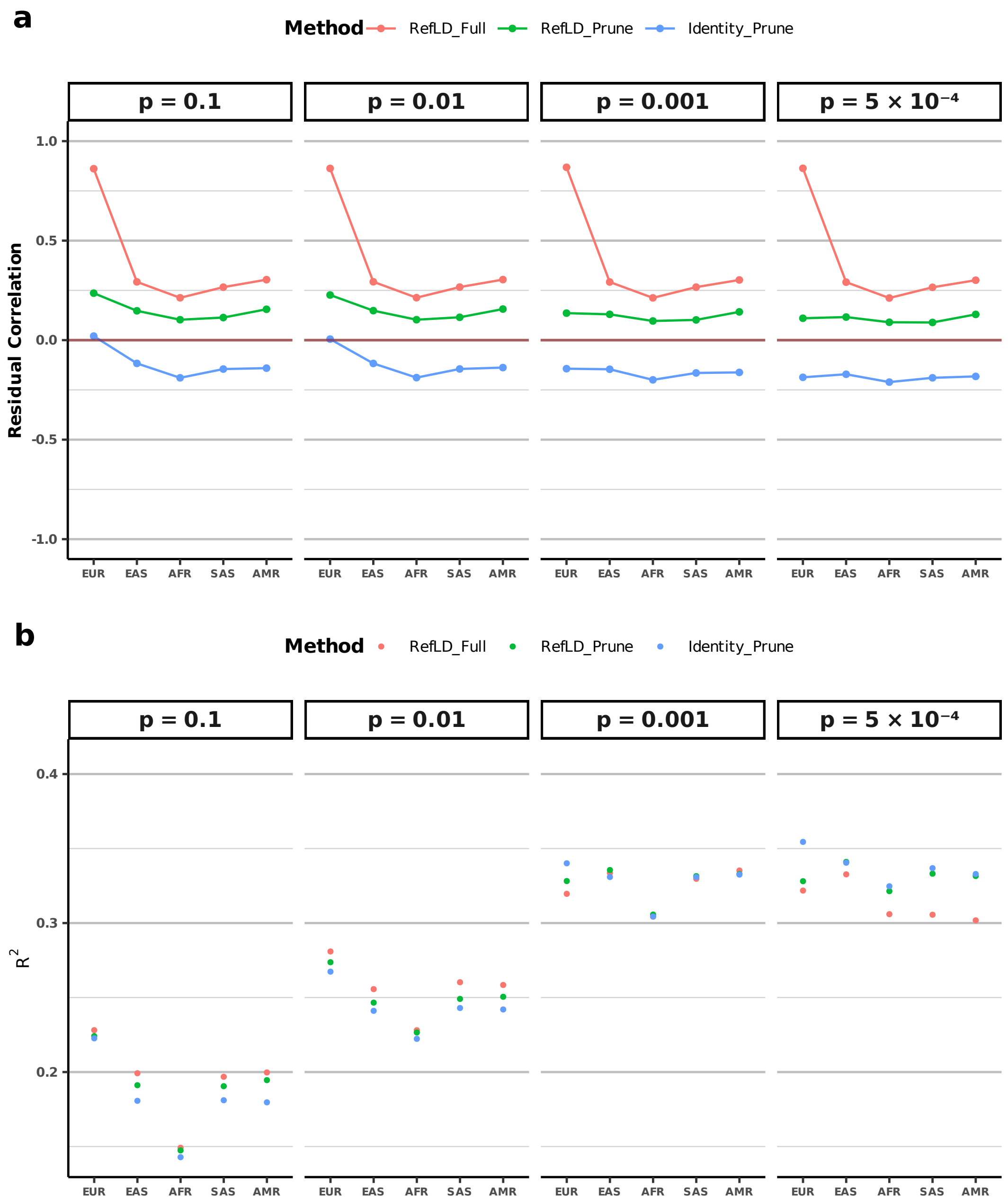

**Figure S3. Relative improvement of MIXPRS using SNP pruning with identity covariance over MIXPRS with LD reference panel using full SNPs across 26 traits without tuning data (UKBB).** **a–d**, Relative improvement of the SNP pruning strategy compared to the full SNPs strategy evaluated across 26 traits in four non-European populations (**a** EAS; **b** AFR; **c** SAS; and **d** AMR) within UKBB, without using individual-level tuning data. Relative performance was calculated as $R^2_{\text{Prune}}/R^2_{\text{Full}} - 1$ for quantitative traits and $\text{AUC}_{\text{Prune}}/\text{AUC}_{\text{Full}} - 1$ for binary traits. Pink bars indicate positive improvement (pruning strategy outperforms full SNPs strategy), and blue bars indicate negative improvement (full SNPs strategy outperforms pruning strategy).

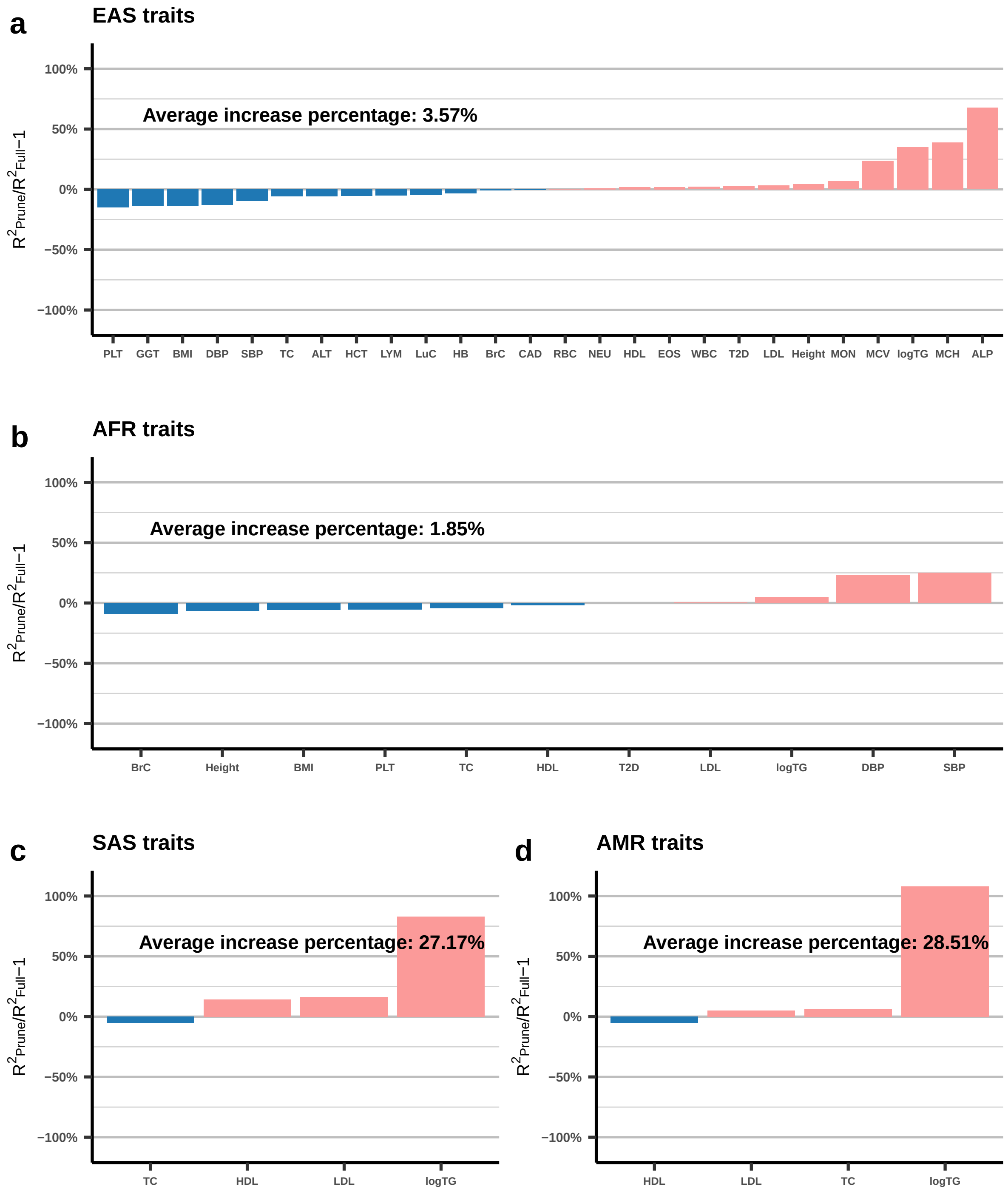

**Figure S4. Relative improvement of MIXPRS using NNLS over MIXPRS using linear regression across 26 traits without tuning data (UKBB). a–d**, Relative improvement of NNLS compared to the linear regression evaluated across 26 traits in four non-European populations (**a** EAS; **b** AFR; **c** SAS; and **d** AMR) within UKBB, without individual-level tuning data. Relative performance was calculated as $R^2_{\text{NNLS}}/R^2_{\text{Linear}} - 1$ for quantitative traits and $\text{AUC}_{\text{NNLS}}/\text{AUC}_{\text{Linear}} - 1$ for binary traits. Pink bars indicate positive improvement (NNLS outperforms linear regression), and blue bars indicate negative improvement (linear regression outperforms NNLS).

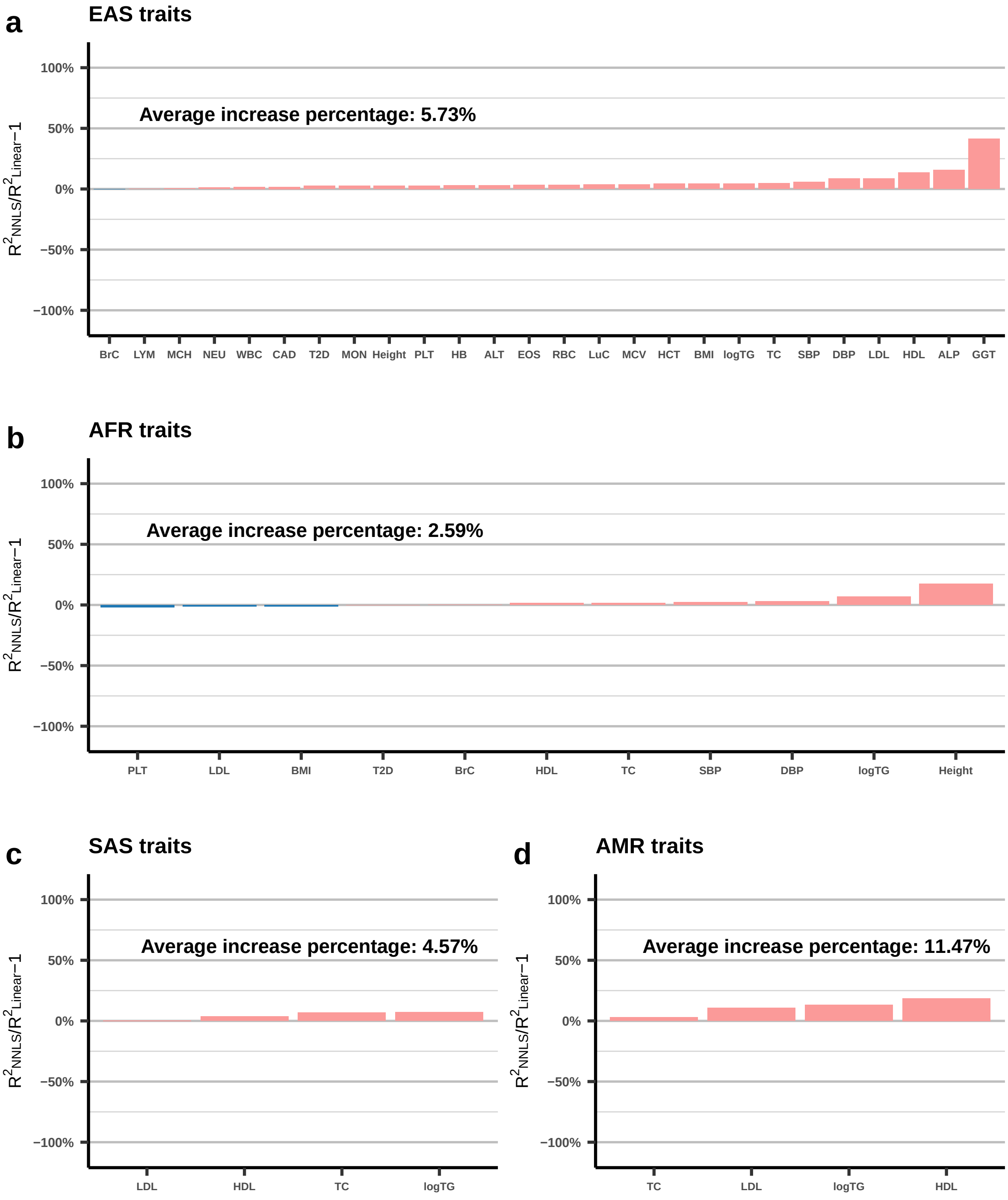

**Figure S5. Prediction accuracy of MIXPRS using SNP pruning with identity covariance versus SNP pruning with LD reference panel across 26 traits without tuning data (UKBB). a–b**, Prediction accuracy of MIXPRS with identity covariance and MIXPRS with LD reference panel were evaluated across (**a**) 22 quantitative traits and (**b**) four binary traits in four non-European populations (EAS, AFR, SAS, and AMR) within UKBB, without individual-level tuning data. Performance metrics were $R^2$ for quantitative traits and AUC for binary traits. Results are displayed as violin plots, with each dot representing the prediction accuracy for an individual trait, and the mean accuracy across traits indicated by a black crossbar within each violin.

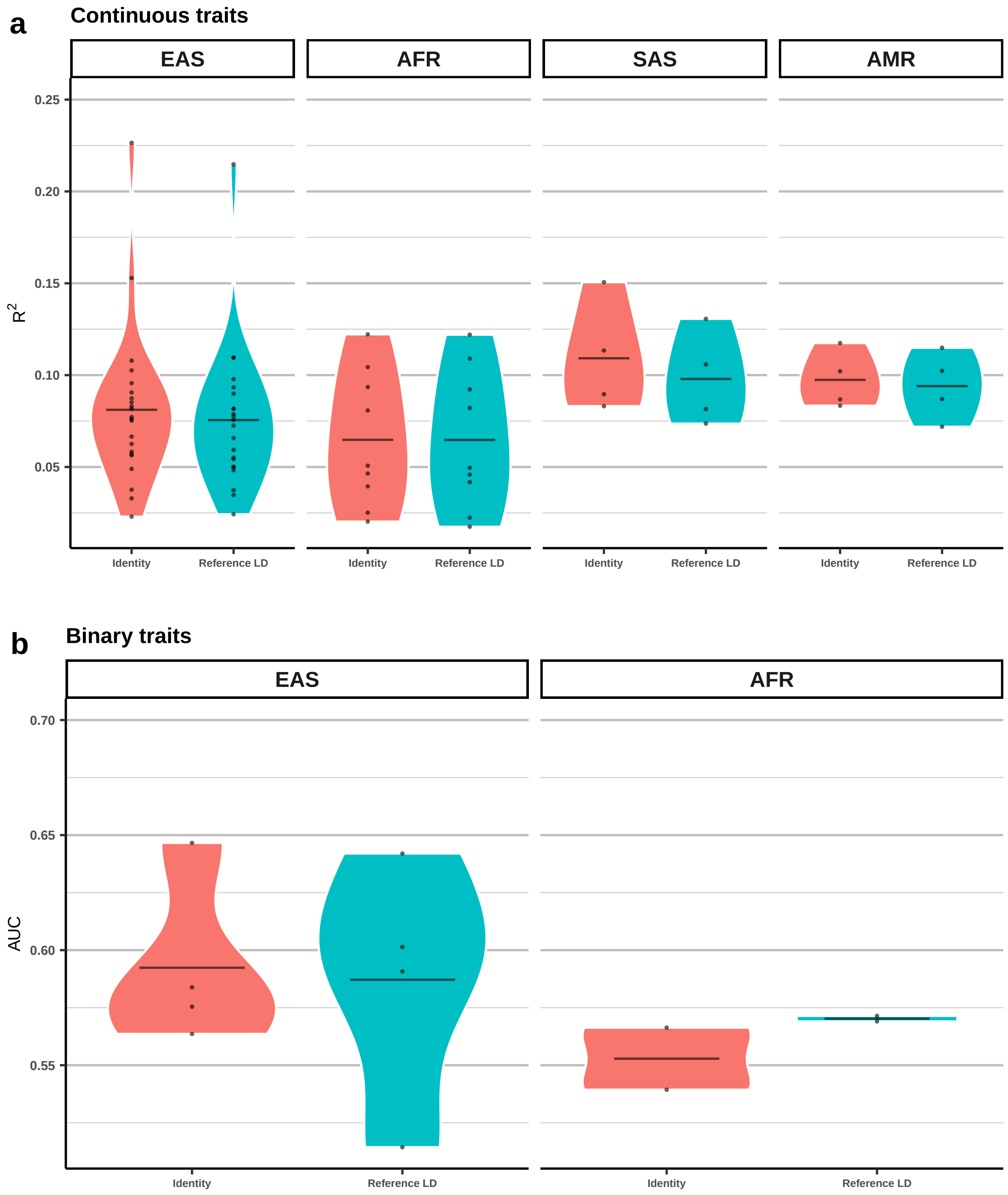

**Figure S6. Prediction accuracy of MIXPRS with different SNP pruning lists for 26 traits without tuning data (UKBB). a–b**, Prediction accuracy of MIXPRS with different SNP pruning lists were evaluated across **a** 22 quantitative traits and **b** four binary traits in four non-European populations (EAS, AFR, SAS, and AMR) within UKBB, without individual-level tuning data. Performance metrics used were $R^2$ for quantitative traits and AUC for binary traits. Results are presented as violin plots, with each dot representing the prediction accuracy for an individual trait, and the mean accuracy across traits indicated by a black crossbar within each violin.

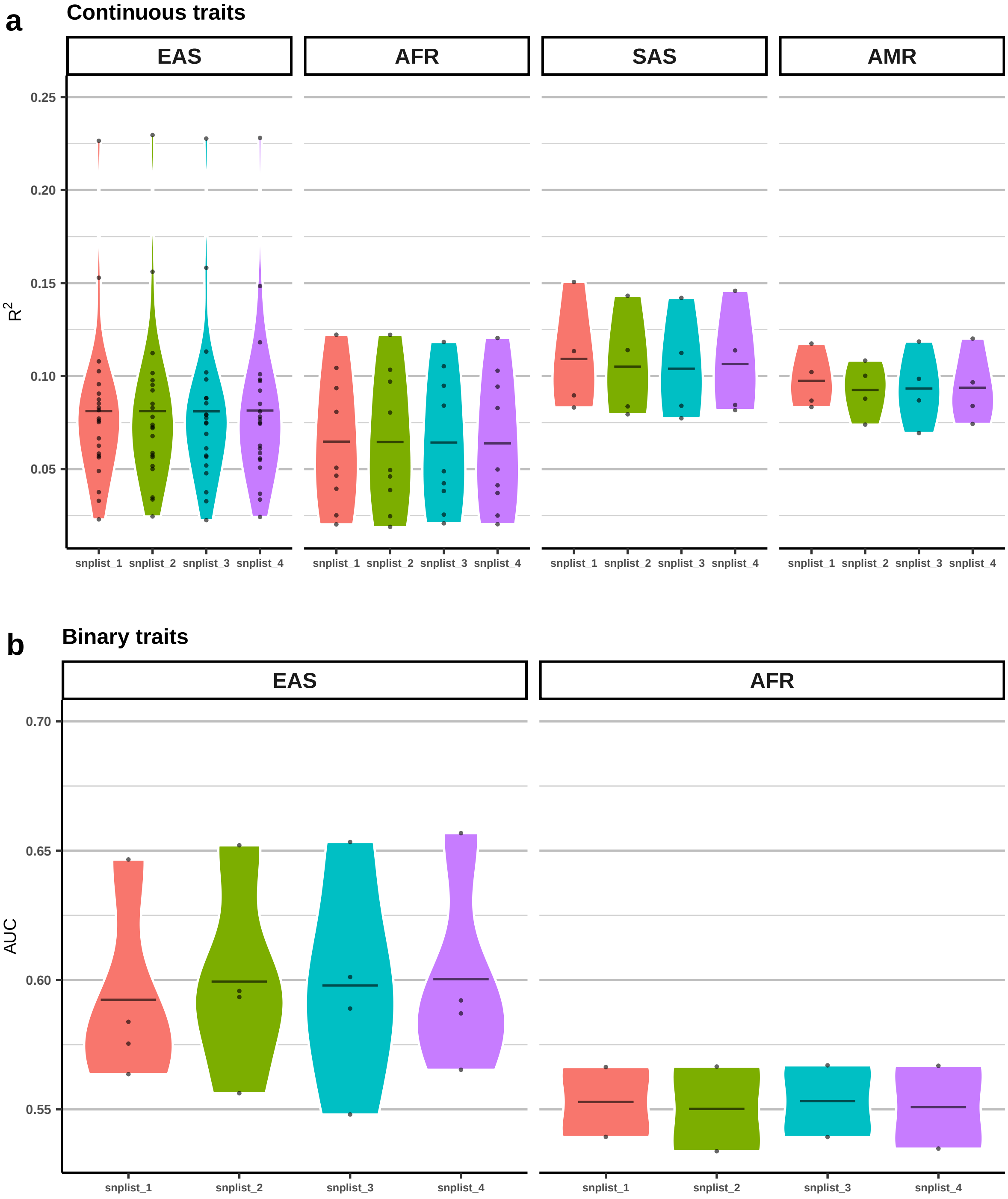

**Figure S7. Prediction accuracy of MIXPRS and its component methods (JointPRS-auto and SDPRX) across 26 traits without tuning data (UKBB). a–f**, Prediction accuracy of MIXPRS and its component methods (JointPRS-auto and SDPRX) were evaluated across 26 traits in four non-European populations within UKBB, without individual-level tuning data. Panels depict results for continuous traits in **a** EAS, **b** AFR, **c** SAS, and **d** AMR, and binary traits in **e** EAS and **f** AFR. Evaluation metrics were $R^2$ for quantitative traits and AUC for binary traits. Results are presented as dot plots, with each dot representing prediction accuracy for an individual trait for each method.

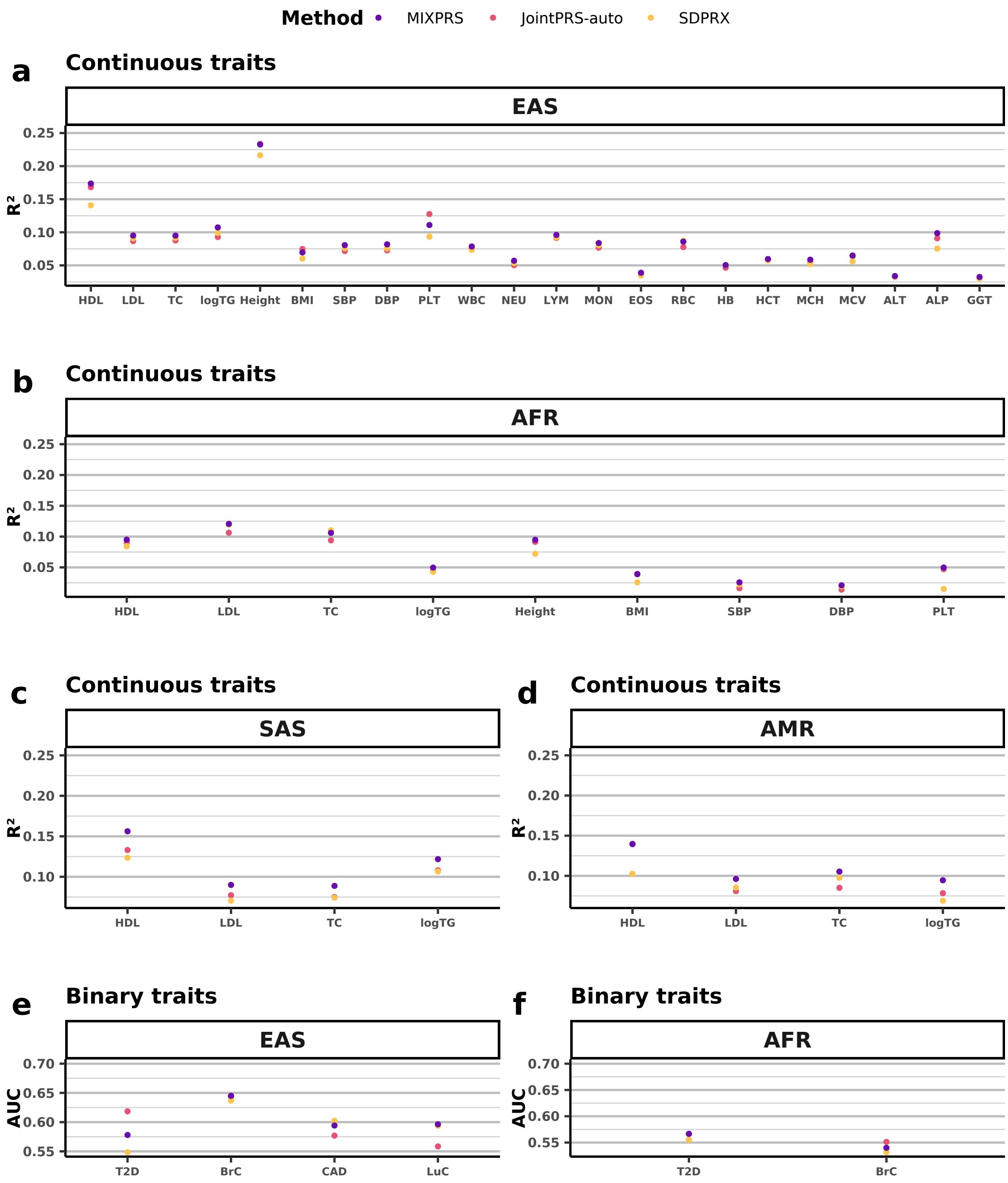

**Figure S8. Prediction accuracy of MIXPRS and PUMAS-EN across four lipid traits without tuning data (UKBB). a–c**, Prediction accuracy of MIXPRS, PUMAS-EN, and PUMAS-EN_paper was evaluated for four lipid traits in three non-European populations (**a** EAS, **b** AFR, and **c** SAS) within UKBB, without individual-level tuning data. Performance was assessed using $R^2$ for these quantitative traits, with results displayed as bar plots. The best-performing and second-best-performing methods are indicated by two stars and one star, respectively, above the corresponding bars.

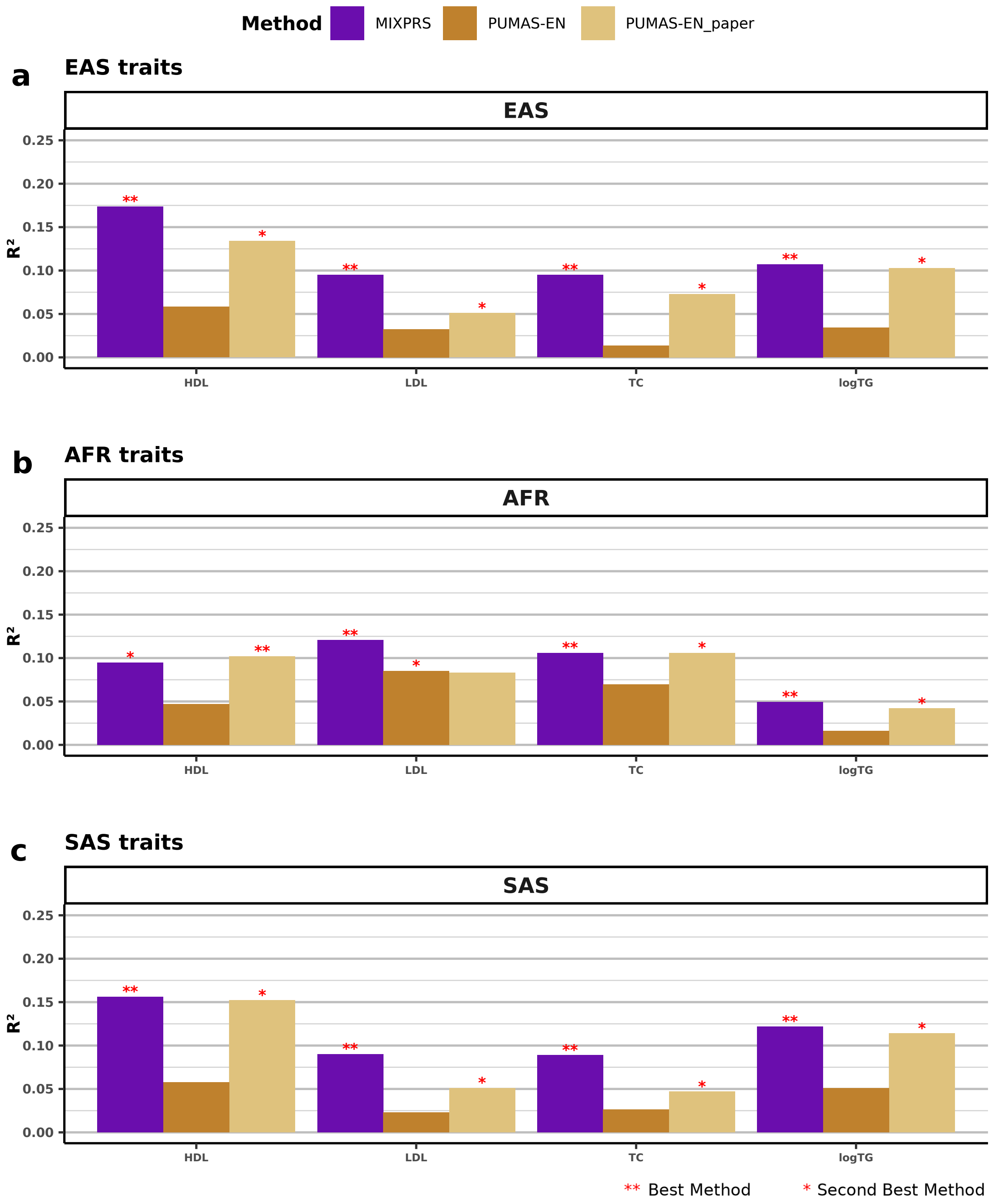

**Figure S9. Prediction accuracy of MIXPRS, IndPRS and SDPRX across nine traits when tuning and testing data are from different cohorts (UKBB and AoU). a–b**, Prediction accuracy of MIXPRS, IndPRS, and SDPRX were evaluated across nine traits in two non-European populations (**a** AFR; **b** AMR) using tuning data from UKBB and testing data from AoU. Evaluation metrics were $R^2$ for quantitative traits and AUC for binary traits. Results are presented as dot plots, with each dot representing prediction accuracy for an individual trait for each method.

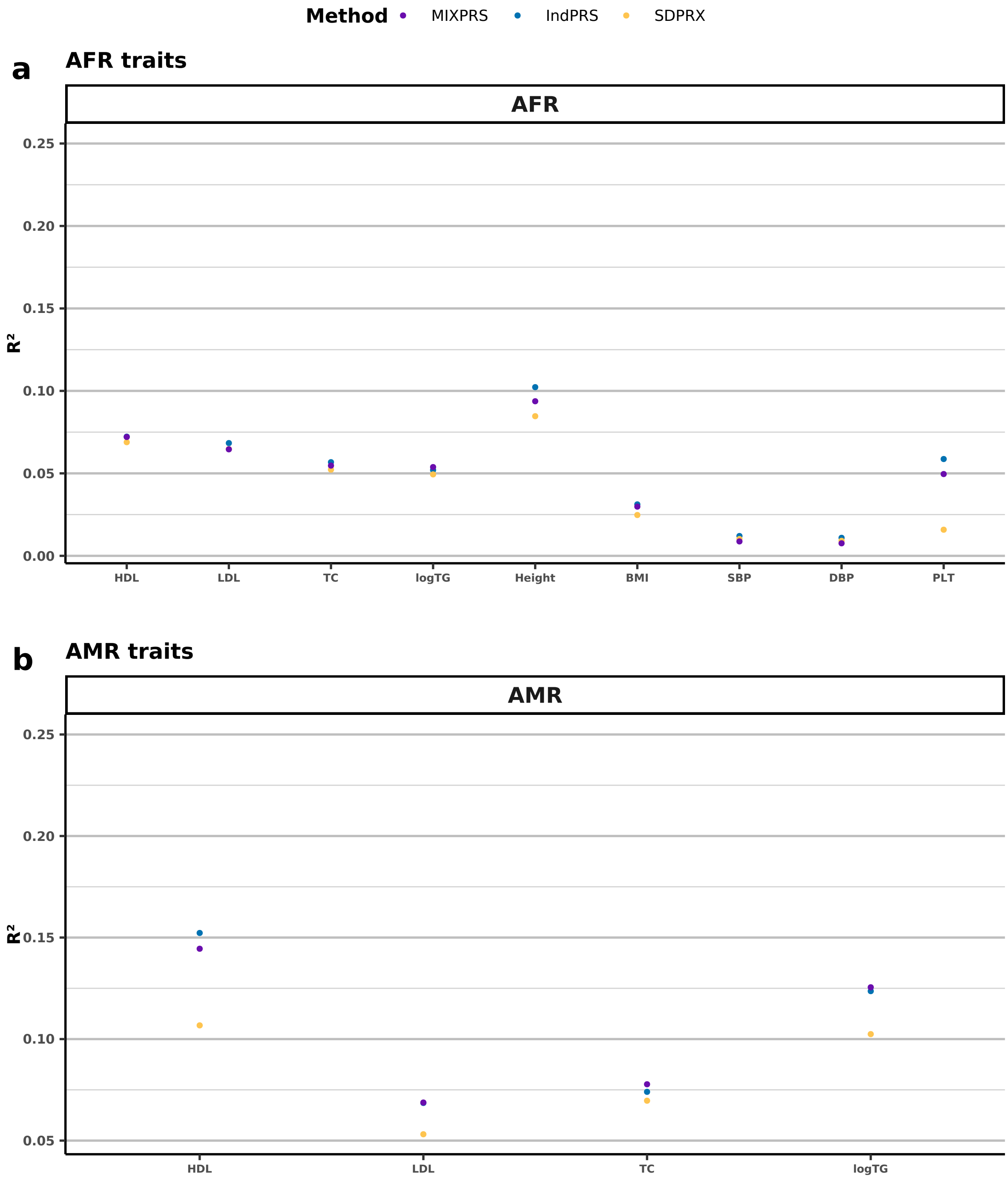

### 2 Supplemental Tables

**Table S1. Summary of sample sizes in GWAS summary statistics for 26 traits *.**

| **Continuous traits** | | **GWAS sample size (EUR; EAS; AFR; SAS; AMR)** |
|---|---|---|
| HDL | HDL-cholesterol | 885,546 [1]; 116,404 [1]; 90,804 [1]; 33,953 [1]; 47,276 [1] |
| LDL | LDL-cholesterol | 840,012 [1]; 79,693 [1]; 87,759 [1]; 33,658 [1]; 33,989 [1] |
| TC | Total cholesterol | 929,739 [1]; 144,579 [1]; 92,554 [1]; 34,135 [1]; 48,055 [1] |
| logTG | Triglycerides | 860,679 [1]; 81,071 [1]; 89,467 [1]; 34,023 [1]; 37,273 [1] |
| Height | Height | 252,357 [2]; 159,095 [3]; 49,781 [4]; N/A; N/A |
| BMI | Body mass index | 233,787 [5]; 158,284 [6]; 49,335 [4]; N/A; N/A |
| SBP | Systolic blood pressure | 728,893 [7]; 179,000 [8]; 35,433 [4]; N/A; N/A |
| DBP | Diastolic blood pressure | 746,038 [7]; 179,000 [8]; 35,433 [4]; N/A; N/A |
| PLT | Platelet | 539,667 [9]; 179,000 [8]; 29,328 [4]; N/A; N/A |
| WBC | White blood cell | 559,083 [9]; 179,000 [8]; N/A; N/A; N/A |
| NEU | Neutrophil | 517,889 [9]; 179,000 [8]; N/A; N/A; N/A |
| LYM | Lymphocyte | 523,524 [9]; 179,000 [8]; N/A; N/A; N/A |
| MON | Monocyte | 520,195 [9]; 179,000 [8]; N/A; N/A; N/A |
| EOS | Eosinophil | 473,152 [9]; 179,000 [8]; N/A; N/A; N/A |
| RBC | Red blood cell | 542,043 [9]; 179,000 [8]; N/A; N/A; N/A |
| HCT | Hematocrit | 559,099 [9]; 179,000 [8]; N/A; N/A; N/A |
| MCH | Mean corpuscular hemoglobin | 483,664 [9]; 179,000 [8]; N/A; N/A; N/A |
| MCV | Mean corpuscular volume | 540,967 [9]; 179,000 [8]; N/A; N/A; N/A |
| HB | Hemoglobin | 408,112 [9]; 179,000 [8]; N/A; N/A; N/A |
| ALT | Alanine aminotransferase | 437,267 [10]; 179,000 [8]; N/A; N/A; N/A |
| ALP | Alkaline phosphatase | 437,267 [10]; 179,000 [8]; N/A; N/A; N/A |
| GGT | $\gamma$-glutamyl transpeptidase | 437,267 [10]; 179,000 [8]; N/A; N/A; N/A |
| **Binary traits** | | **GWAS cases/controls sample size (EUR; EAS; AFR)** |
| T2D | Type 2 diabetes | 26,676/132,532 [11]; 36,614/155,150 [12]; 14,042/31,683 [4] |
| BrC | Breast cancer | 133,384/113,789 [13]; 14,068/13,104 [14, 8]; 4832/3020 [14] |
| CAD | Coronary artery disease | 22,233/64,762 [15]; 29,319/183,134 [16]; N/A |
| LuC | Lung cancer | 29,266/56,450 [17]; 4,050/208,403 [16]; N/A |

* Note: All non-European GWAS summary statistics exclude individuals from UKBB or AoU cohorts.

**Table S2. Summary of SNP number in GWAS summary statistics for 26 traits.**

| Continuous traits | | | | | | |
|---|---|---|---|---|---|---|
| Trait | | GWAS SNP number | | | | |
| | | EUR | EAS | AFR | SAS | AMR |
| HDL | HDL-cholesterol | 800,281 | 735,249 | 827,727 | 1,085,452 | 1,107,923 |
| LDL | LDL-cholesterol | 800,283 | 797,861 | 827,727 | 1,088,264 | 1,104,517 |
| TC | Total cholesterol | 800,281 | 461,893 | 827,727 | 1,085,270 | 1,111,066 |
| logTG | Triglycerides | 800,286 | 797,898 | 827,727 | 1,088,215 | 1,109,126 |
| Height | Height | 724,431 | 790,675 | 827,738 | N/A | N/A |
| BMI | Body mass index | 725,221 | 782,322 | 827,738 | N/A | N/A |
| SBP | Systolic blood pressure | 797,661 | 747,306 | 827,738 | N/A | N/A |
| DBP | Diastolic blood pressure | 798,292 | 747,306 | 827,738 | N/A | N/A |
| PLT | Platelet | 800,321 | 747,306 | 827,738 | N/A | N/A |
| WBC | White blood cell | 800,320 | 747,306 | N/A | N/A | N/A |
| NEU | Neutrophil | 800,319 | 747,306 | N/A | N/A | N/A |
| LYM | Lymphocyte | 800,320 | 747,306 | N/A | N/A | N/A |
| MON | Monocyte | 800,320 | 747,306 | N/A | N/A | N/A |
| EOS | Eosinophil | 800,319 | 747,306 | N/A | N/A | N/A |
| RBC | Red blood cell | 800,318 | 747,306 | N/A | N/A | N/A |
| HCT | Hematocrit | 800,319 | 747,306 | N/A | N/A | N/A |
| MCH | Mean corpuscular hemoglobin | 800,319 | 747,306 | N/A | N/A | N/A |
| MCV | Mean corpuscular volume | 800,317 | 747,306 | N/A | N/A | N/A |
| HB | Hemoglobin | 800,322 | 747,306 | N/A | N/A | N/A |
| ALT | Alanine aminotransferase | 787,866 | 747,306 | N/A | N/A | N/A |
| ALP | Alkaline phosphatase | 799,800 | 747,306 | N/A | N/A | N/A |
| GGT | $\gamma$-glutamyl transpeptidase | 799,800 | 747,306 | N/A | N/A | N/A |

| Binary traits | | | | |
|---|---|---|---|---|
| Trait | | GWAS SNP number | | |
| | | EUR | EAS | AFR |
| T2D | Type 2 diabetes | 800,320 | 799,544 | 827,738 |
| BrC | Breast cancer | 800,245 | 747,306 | 837,774 |
| CAD | Coronary artery disease | 653,825 | 786,215 | N/A |
| LuC | Lung cancer | 756,305 | 786,215 | N/A |

**Table S3. Summary of sample sizes in UKBB for 26 traits.**

**Continuous traits**

| Trait | | UKBB sample size | | | | UKBB Field |
|---|---|---|---|---|---|---|
| | | **EAS** | **AFR** | **SAS** | **AMR** | |
| HDL | HDL-cholesterol | 1,812 | 5,927 | 6,784 | 561 | 30760 |
| LDL | LDL-cholesterol | 1,911 | 6,171 | 7,062 | 583 | 30780 |
| TC | Total cholesterol | 1,912 | 6,183 | 7,080 | 583 | 30690 |
| logTG | Triglycerides | 1,993 | 6,400 | 7,444 | 607 | 30870 |
| Height | Height | 2,080 | 6,574 | N/A | N/A | 50 |
| BMI | Body mass index | 2,077 | 6,713 | N/A | N/A | 21001 |
| SBP | Systolic blood pressure | 2,001 | 6,688 | N/A | N/A | 93; 4080 |
| DBP | Diastolic blood pressure | 2,001 | 6,574 | N/A | N/A | 94; 4090 |
| PLT | Platelet | 2,027 | 6,445 | N/A | N/A | 30080 |
| WBC | White blood cell | 2,027 | N/A | N/A | N/A | 30000 |
| NEU | Neutrophil | 2,025 | N/A | N/A | N/A | 30140 |
| LYM | Lymphocyte | 2,025 | N/A | N/A | N/A | 30120 |
| MON | Monocyte | 2,025 | N/A | N/A | N/A | 30130 |
| EOS | Eosinophil | 2,025 | N/A | N/A | N/A | 30150 |
| RBC | Red blood cell | 2,027 | N/A | N/A | N/A | 30010 |
| HCT | Hematocrit | 2,027 | N/A | N/A | N/A | 30030 |
| MCH | Mean corpuscular hemoglobin | 2,027 | N/A | N/A | N/A | 30050 |
| MCV | Mean corpuscular volume | 2,027 | N/A | N/A | N/A | 30040 |
| HB | Hemoglobin | 2,027 | N/A | N/A | N/A | 30020 |
| ALT | Alanine aminotransferase | 1,993 | N/A | N/A | N/A | 30620 |
| ALP | Alkaline phosphatase | 1,994 | N/A | N/A | N/A | 30610 |
| GGT | $\gamma$-glutamyl transpeptidase | 1,991 | N/A | N/A | N/A | 30730 |

**Binary traits**

| Trait | | UKBB sample size | | | |
|---|---|---|---|---|---|
| | | **EAS** | | **AFR** | |
| | | **case** | **control** | **case** | **control** |
| T2D | Type 2 diabetes | 203 | 1,887 | 1,292 | 5,535 |
| BrC | Breast cancer | 89 | 2,001 | 173 | 6,654 |
| CAD | Coronary artery disease | 56 | 2,034 | N/A | N/A |
| LuC | Lung cancer | 21 | 2,069 | N/A | N/A |

**Table S4. Summary of sample sizes in AoU for nine traits.**

| Continuous traits | | | | |
|---|---|---|---|---|
| Trait | | AoU sample size | | AoU Concept Id |
| | | AFR | AMR | |
| HDL | HDL-cholesterol | 12822 | 7004 | 3007070 |
| LDL | LDL-cholesterol | 9359 | 5457 | 3028288 |
| TC | Total cholesterol | 12429 | 7016 | 3027114 |
| logTG | Triglycerides | 12395 | 7020 | 3022192 |
| Trait | | AoU sample size | | AoU Concept Id |
| | | AFR | AMR | |
| Height | Height | 51790 | N/A | 903133 |
| BMI | Body mass index | 50747 | N/A | 903124 |
| SBP | Systolic blood pressure | 51342 | N/A | 903118 |
| DBP | Diastolic blood pressure | 51395 | N/A | 903115 |
| PLT | Platelet | 24517 | N/A | 37037425 |

**Table S5. Simulation results of eight methods for mean derivation of $R^2$ across five replications.**

| p | Method | EAS | | AFR | | SAS | | AMR | |
|---|---|---|---|---|---|---|---|---|---|
| | | **25K** | **90K** | **25K** | **90K** | **25K** | **90K** | **25K** | **90K** |
| p = 0.1 | MIXPRS | 0.125 | 0.182 | 0.081 | 0.139 | 0.128 | 0.184 | 0.131 | 0.183 |
| | JointPRS | 0.115 | 0.164 | 0.075 | 0.124 | 0.119 | 0.169 | 0.121 | 0.168 |
| | XPASS | 0.097 | 0.127 | 0.067 | 0.090 | 0.106 | 0.140 | 0.095 | 0.082 |
| | SDPRX | 0.112 | 0.155 | 0.073 | 0.115 | 0.114 | 0.154 | 0.117 | 0.156 |
| | PRS-CSx | 0.116 | 0.116 | 0.081 | 0.081 | 0.123 | 0.123 | 0.126 | 0.126 |
| | MUSSEL | 0.094 | 0.094 | 0.056 | 0.056 | 0.094 | 0.094 | 0.087 | 0.087 |
| | PROSPER | 0.087 | 0.087 | 0.045 | 0.045 | 0.097 | 0.097 | 0.090 | 0.090 |
| | BridgePRS | 0.090 | 0.090 | 0.061 | 0.061 | 0.094 | 0.094 | 0.099 | 0.099 |
| p = 0.01 | MIXPRS | 0.183 | 0.242 | 0.150 | 0.210 | 0.186 | 0.240 | 0.191 | 0.244 |
| | JointPRS | 0.179 | 0.227 | 0.145 | 0.197 | 0.185 | 0.231 | 0.188 | 0.232 |
| | XPASS | 0.065 | 0.102 | 0.043 | 0.066 | 0.074 | 0.120 | 0.055 | 0.039 |
| | SDPRX | 0.162 | 0.213 | 0.132 | 0.174 | 0.163 | 0.204 | 0.170 | 0.212 |
| | PRS-CSx | 0.154 | 0.154 | 0.119 | 0.119 | 0.160 | 0.160 | 0.163 | 0.163 |
| | MUSSEL | 0.145 | 0.145 | 0.125 | 0.125 | 0.143 | 0.143 | 0.153 | 0.153 |
| | PROSPER | 0.126 | 0.126 | 0.079 | 0.079 | 0.135 | 0.135 | 0.119 | 0.119 |
| | BridgePRS | 0.101 | 0.101 | 0.070 | 0.070 | 0.115 | 0.115 | 0.119 | 0.119 |
| p = 0.001 | MIXPRS | 0.284 | 0.316 | 0.260 | 0.296 | 0.285 | 0.317 | 0.283 | 0.312 |
| | JointPRS | 0.271 | 0.298 | 0.245 | 0.277 | 0.273 | 0.300 | 0.269 | 0.296 |
| | XPASS | 0.075 | 0.238 | 0.074 | 0.230 | 0.076 | 0.235 | 0.074 | 0.236 |
| | SDPRX | 0.272 | 0.313 | 0.250 | 0.299 | 0.274 | 0.313 | 0.273 | 0.303 |
| | PRS-CSx | 0.259 | 0.259 | 0.231 | 0.231 | 0.265 | 0.265 | 0.260 | 0.260 |
| | MUSSEL | 0.231 | 0.231 | 0.229 | 0.229 | 0.175 | 0.175 | 0.216 | 0.216 |
| | PROSPER | 0.227 | 0.227 | 0.199 | 0.199 | 0.235 | 0.235 | 0.231 | 0.231 |
| | BridgePRS | 0.139 | 0.139 | 0.106 | 0.106 | 0.168 | 0.168 | 0.166 | 0.166 |
| p = $5 \times 10^{-4}$ | MIXPRS | 0.299 | 0.332 | 0.271 | 0.308 | 0.299 | 0.324 | 0.301 | 0.324 |
| | JointPRS | 0.295 | 0.318 | 0.260 | 0.288 | 0.291 | 0.312 | 0.292 | 0.314 |
| | XPASS | 0.153 | 0.286 | 0.136 | 0.271 | 0.149 | 0.269 | 0.159 | 0.291 |
| | SDPRX | 0.298 | 0.332 | 0.266 | 0.308 | 0.290 | 0.322 | 0.297 | 0.325 |
| | PRS-CSx | 0.289 | 0.289 | 0.253 | 0.253 | 0.285 | 0.285 | 0.288 | 0.288 |
| | MUSSEL | 0.227 | 0.227 | 0.259 | 0.259 | 0.208 | 0.208 | 0.226 | 0.226 |
| | PROSPER | 0.261 | 0.261 | 0.228 | 0.228 | 0.257 | 0.257 | 0.263 | 0.263 |
| | BridgePRS | 0.176 | 0.176 | 0.140 | 0.140 | 0.192 | 0.192 | 0.197 | 0.197 |

**Table S6. Simulation results of MIXPRS and its component methods (JointPRS-auto and SDPRX) for mean derivation of $R^2$ across five replications.**

| p | Method | EUR | EAS | AFR | SAS | AMR |
|---|---|---|---|---|---|---|
| | | **100K** | **100K** | **100K** | **100K** | **100K** |
| p = 0.1 | MIXPRS | 0.227 | 0.196 | 0.148 | 0.195 | 0.200 |
| | JointPRS-auto | 0.215 | 0.195 | 0.143 | 0.193 | 0.197 |
| | SDPRX | 0.206 | 0.167 | 0.123 | 0.167 | 0.171 |
| p = 0.01 | MIXPRS | 0.275 | 0.253 | 0.226 | 0.256 | 0.254 |
| | JointPRS-auto | 0.282 | 0.252 | 0.224 | 0.259 | 0.256 |
| | SDPRX | 0.262 | 0.224 | 0.191 | 0.219 | 0.224 |
| p = 0.001 | MIXPRS | 0.340 | 0.325 | 0.298 | 0.327 | 0.328 |
| | JointPRS-auto | 0.336 | 0.318 | 0.287 | 0.321 | 0.319 |
| | SDPRX | 0.336 | 0.329 | 0.301 | 0.323 | 0.327 |
| p = $5 \times 10^{-4}$ | MIXPRS | 0.354 | 0.340 | 0.323 | 0.336 | 0.331 |
| | JointPRS-auto | 0.353 | 0.338 | 0.310 | 0.333 | 0.332 |
| | SDPRX | 0.347 | 0.341 | 0.321 | 0.334 | 0.329 |

**Table S7. Simulation results of MIXPRS with different GWAS subsampling strategies for mean derivation of residual correlations across five replications.**

| p | Strategy | EUR | EAS | AFR | SAS | AMR |
|---|---|---|---|---|---|---|
| p = 0.1 | RefLD_Full | 0.861 | 0.293 | 0.213 | 0.266 | 0.304 |
| | RefLD_Prune | 0.236 | 0.147 | 0.103 | 0.114 | 0.155 |
| | Identity_Prune | 0.020 | -0.117 | -0.189 | -0.145 | -0.141 |
| p = 0.01 | RefLD_Full | 0.863 | 0.293 | 0.213 | 0.267 | 0.304 |
| | RefLD_Prune | 0.227 | 0.148 | 0.103 | 0.115 | 0.156 |
| | Identity_Prune | 0.005 | -0.117 | -0.188 | -0.145 | -0.138 |
| p = 0.001 | RefLD_Full | 0.869 | 0.292 | 0.212 | 0.266 | 0.302 |
| | RefLD_Prune | 0.136 | 0.130 | 0.096 | 0.102 | 0.142 |
| | Identity_Prune | -0.143 | -0.146 | -0.200 | -0.165 | -0.162 |
| $p = 5 \times 10^{-4}$ | RefLD_Full | 0.864 | 0.291 | 0.212 | 0.266 | 0.301 |
| | RefLD_Prune | 0.110 | 0.116 | 0.090 | 0.089 | 0.129 |
| | Identity_Prune | -0.187 | -0.171 | -0.211 | -0.189 | -0.182 |

**Table S8. Simulation results of MIXPRS with different GWAS subsampling strategies for mean derivation of $R^2$ across five replications.**

| p | Strategy | EUR | EAS | AFR | SAS | AMR |
|---|---|---|---|---|---|---|
| p = 0.1 | RefLD_Full | 0.228 | 0.199 | 0.149 | 0.197 | 0.200 |
| | RefLD_Prune | 0.224 | 0.191 | 0.147 | 0.191 | 0.195 |
| | Identity_Prune | 0.223 | 0.181 | 0.143 | 0.181 | 0.180 |
| p = 0.01 | RefLD_Full | 0.281 | 0.256 | 0.228 | 0.260 | 0.258 |
| | RefLD_Prune | 0.274 | 0.247 | 0.227 | 0.249 | 0.251 |
| | Identity_Prune | 0.267 | 0.241 | 0.222 | 0.243 | 0.242 |
| p = 0.001 | RefLD_Full | 0.320 | 0.334 | 0.304 | 0.330 | 0.335 |
| | RefLD_Prune | 0.328 | 0.336 | 0.306 | 0.332 | 0.333 |
| | Identity_Prune | 0.340 | 0.331 | 0.304 | 0.331 | 0.333 |
| p = $5 \times 10^{-4}$ | RefLD_Full | 0.322 | 0.333 | 0.306 | 0.306 | 0.302 |
| | RefLD_Prune | 0.328 | 0.341 | 0.321 | 0.333 | 0.332 |
| | Identity_Prune | 0.354 | 0.340 | 0.325 | 0.337 | 0.333 |

**Table S9. Real data results of MIXPRS with different strategies for $R^2$ or AUC of 26 traits without tuning data (UKBB).**

| Pop | Trait | Prune_Linear | Prune_NNLS | Full_Linear | Pop | Trait | Prune_Linear | Prune_NNLS | Full_Linear |
|---|---|---|---|---|---|---|---|---|---|
| EAS | HDL | 0.153 | 0.174 | 0.150 | AFR | HDL | 0.094 | 0.095 | 0.096 |
| | LDL | 0.087 | 0.095 | 0.085 | | LDL | 0.122 | 0.121 | 0.121 |
| | TC | 0.091 | 0.095 | 0.096 | | TC | 0.104 | 0.106 | 0.109 |
| | logTG | 0.103 | 0.107 | 0.076 | | logTG | 0.046 | 0.050 | 0.044 |
| | Height | 0.226 | 0.233 | 0.217 | | Height | 0.081 | 0.095 | 0.086 |
| | BMI | 0.067 | 0.070 | 0.077 | | BMI | 0.039 | 0.039 | 0.042 |
| | SBP | 0.076 | 0.081 | 0.084 | | SBP | 0.025 | 0.026 | 0.020 |
| | DBP | 0.075 | 0.082 | 0.086 | | DBP | 0.020 | 0.021 | 0.016 |
| | PLT | 0.108 | 0.111 | 0.127 | | PLT | 0.051 | 0.050 | 0.054 |
| | WBC | 0.077 | 0.079 | 0.076 | | T2D | 0.566 | 0.567 | 0.565 |
| | NEU | 0.056 | 0.057 | 0.056 | | BrC | 0.539 | 0.540 | 0.592 |
| | LYM | 0.096 | 0.096 | 0.101 | SAS | HDL | 0.151 | 0.156 | 0.132 |
| | MON | 0.082 | 0.084 | 0.077 | | LDL | 0.090 | 0.090 | 0.077 |
| | EOS | 0.038 | 0.039 | 0.037 | | TC | 0.083 | 0.089 | 0.087 |
| | RBC | 0.083 | 0.086 | 0.083 | | logTG | 0.113 | 0.122 | 0.062 |
| | HB | 0.049 | 0.050 | 0.051 | AMR | HDL | 0.117 | 0.139 | 0.124 |
| | HCT | 0.057 | 0.060 | 0.060 | | LDL | 0.087 | 0.096 | 0.083 |
| | MCH | 0.058 | 0.059 | 0.042 | | TC | 0.102 | 0.105 | 0.096 |
| | MCV | 0.063 | 0.065 | 0.051 | | logTG | 0.083 | 0.094 | 0.040 |
| | ALT | 0.033 | 0.034 | 0.035 | | | | | |
| | ALP | 0.085 | 0.099 | 0.051 | | | | | |
| | GGT | 0.023 | 0.032 | 0.027 | | | | | |
| | T2D | 0.564 | 0.578 | 0.547 | | | | | |
| | BrC | 0.647 | 0.645 | 0.653 | | | | | |
| | CAD | 0.584 | 0.594 | 0.588 | | | | | |
| | LuC | 0.575 | 0.596 | 0.605 | | | | | |

**Table S10. Real data results of MIXPRS with different SNP covariance structures for $R^2$ or AUC of 26 traits without tuning data (UKBB).**

| Pop | Trait | Identity | Reference LD | Pop | Trait | Identity | Reference LD |
|---|---|---|---|---|---|---|---|
| EAS | HDL | 0.153 | 0.110 | AFR | HDL | 0.094 | 0.092 |
| | LDL | 0.087 | 0.090 | | LDL | 0.122 | 0.122 |
| | TC | 0.091 | 0.093 | | TC | 0.104 | 0.109 |
| | logTG | 0.103 | 0.073 | | logTG | 0.046 | 0.046 |
| | Height | 0.226 | 0.215 | | Height | 0.081 | 0.082 |
| | BMI | 0.067 | 0.066 | | BMI | 0.039 | 0.042 |
| | SBP | 0.076 | 0.082 | | SBP | 0.025 | 0.022 |
| | DBP | 0.075 | 0.082 | | DBP | 0.020 | 0.017 |
| | PLT | 0.108 | 0.110 | | PLT | 0.051 | 0.050 |
| | WBC | 0.077 | 0.077 | | T2D | 0.566 | 0.569 |
| | NEU | 0.056 | 0.054 | | BrC | 0.539 | 0.571 |
| | LYM | 0.096 | 0.098 | SAS | HDL | 0.151 | 0.131 |
| | MON | 0.082 | 0.076 | | LDL | 0.090 | 0.074 |
| | EOS | 0.038 | 0.037 | | TC | 0.083 | 0.082 |
| | RBC | 0.083 | 0.079 | | logTG | 0.113 | 0.106 |
| | HB | 0.049 | 0.050 | AMR | HDL | 0.117 | 0.115 |
| | HCT | 0.057 | 0.059 | | LDL | 0.087 | 0.087 |
| | MCH | 0.058 | 0.050 | | TC | 0.102 | 0.102 |
| | MCV | 0.063 | 0.055 | | logTG | 0.083 | 0.072 |
| | ALT | 0.033 | 0.035 | | | | |
| | ALP | 0.085 | 0.048 | | | | |
| | GGT | 0.023 | 0.024 | | | | |
| | T2D | 0.564 | 0.514 | | | | |
| | BrC | 0.647 | 0.642 | | | | |
| | CAD | 0.584 | 0.591 | | | | |
| | LuC | 0.575 | 0.601 | | | | |

**Table S11. Real data results of MIXPRS with different prune SNP lists structures for $R^2$ or AUC of 26 traits without tuning data (UKBB).**

| Pop | Trait | snplist_1 | snplist_2 | snplist_3 | snplist_4 | Pop | Trait | snplist_1 | snplist_2 | snplist_3 | snplist_4 |
|---|---|---|---|---|---|---|---|---|---|---|---|
| EAS | HDL | 0.153 | 0.156 | 0.158 | 0.148 | AFR | HDL | 0.094 | 0.097 | 0.095 | 0.094 |
| | LDL | 0.087 | 0.095 | 0.088 | 0.085 | | LDL | 0.122 | 0.122 | 0.118 | 0.120 |
| | TC | 0.091 | 0.092 | 0.088 | 0.092 | | TC | 0.104 | 0.103 | 0.105 | 0.103 |
| | logTG | 0.103 | 0.102 | 0.102 | 0.101 | | logTG | 0.046 | 0.046 | 0.042 | 0.041 |
| | Height | 0.226 | 0.230 | 0.228 | 0.228 | | Height | 0.081 | 0.080 | 0.084 | 0.083 |
| | BMI | 0.067 | 0.068 | 0.069 | 0.063 | | BMI | 0.039 | 0.039 | 0.038 | 0.037 |
| | SBP | 0.076 | 0.074 | 0.075 | 0.075 | | SBP | 0.025 | 0.025 | 0.025 | 0.025 |
| | DBP | 0.075 | 0.072 | 0.075 | 0.074 | | DBP | 0.020 | 0.019 | 0.021 | 0.020 |
| | PLT | 0.108 | 0.112 | 0.113 | 0.118 | | PLT | 0.051 | 0.049 | 0.049 | 0.050 |
| | WBC | 0.077 | 0.073 | 0.077 | 0.077 | | T2D | 0.566 | 0.567 | 0.567 | 0.567 |
| | NEU | 0.056 | 0.052 | 0.052 | 0.055 | | BrC | 0.539 | 0.534 | 0.539 | 0.535 |
| | LYM | 0.096 | 0.098 | 0.098 | 0.097 | SAS | HDL | 0.151 | 0.143 | 0.142 | 0.146 |
| | MON | 0.082 | 0.083 | 0.079 | 0.078 | | LDL | 0.090 | 0.079 | 0.077 | 0.084 |
| | EOS | 0.038 | 0.034 | 0.038 | 0.037 | | TC | 0.083 | 0.084 | 0.084 | 0.082 |
| | RBC | 0.083 | 0.085 | 0.079 | 0.081 | | logTG | 0.113 | 0.114 | 0.112 | 0.114 |
| | HB | 0.049 | 0.050 | 0.048 | 0.051 | AMR | HDL | 0.117 | 0.108 | 0.119 | 0.120 |
| | HCT | 0.057 | 0.057 | 0.057 | 0.059 | | LDL | 0.087 | 0.088 | 0.087 | 0.084 |
| | MCH | 0.058 | 0.056 | 0.057 | 0.056 | | TC | 0.102 | 0.100 | 0.098 | 0.097 |
| | MCV | 0.063 | 0.059 | 0.061 | 0.061 | | logTG | 0.083 | 0.074 | 0.069 | 0.074 |
| | ALT | 0.033 | 0.035 | 0.033 | 0.034 | | | | | | |
| | ALP | 0.085 | 0.078 | 0.085 | 0.098 | | | | | | |
| | GGT | 0.023 | 0.025 | 0.023 | 0.024 | | | | | | |
| | T2D | 0.564 | 0.556 | 0.548 | 0.565 | | | | | | |
| | BrC | 0.647 | 0.652 | 0.653 | 0.657 | | | | | | |
| | CAD | 0.584 | 0.593 | 0.589 | 0.587 | | | | | | |
| | LuC | 0.575 | 0.596 | 0.601 | 0.592 | | | | | | |

**Table S12. Real data results of five methods for $R^2$ or AUC of 26 traits in EAS without tuning data (UKBB).**

| Pop | Trait | MIXPRS | JointPRS-auto | PRS-CSx-auto | SDPRX | XPASS |
|---|---|---|---|---|---|---|
| EAS | HDL | 0.174 | 0.168 | 0.144 | 0.141 | 0.131 |
| | LDL | 0.095 | 0.087 | 0.082 | 0.091 | 0.074 |
| | TC | 0.095 | 0.088 | 0.076 | 0.092 | 0.068 |
| | logTG | 0.107 | 0.093 | 0.078 | 0.099 | 0.072 |
| | Height | 0.233 | 0.234 | 0.220 | 0.216 | 0.175 |
| | BMI | 0.070 | 0.075 | 0.059 | 0.060 | 0.060 |
| | SBP | 0.081 | 0.072 | 0.055 | 0.076 | 0.045 |
| | DBP | 0.082 | 0.073 | 0.055 | 0.075 | 0.046 |
| | PLT | 0.111 | 0.128 | 0.106 | 0.093 | 0.095 |
| | WBC | 0.079 | 0.075 | 0.056 | 0.073 | 0.040 |
| | NEU | 0.057 | 0.050 | 0.036 | 0.053 | 0.032 |
| | LYM | 0.096 | 0.091 | 0.073 | 0.092 | 0.049 |
| | MON | 0.084 | 0.077 | 0.069 | 0.080 | 0.047 |
| | EOS | 0.039 | 0.036 | 0.029 | 0.035 | 0.019 |
| | RBC | 0.086 | 0.078 | 0.065 | 0.087 | 0.062 |
| | HB | 0.050 | 0.046 | 0.040 | 0.050 | 0.021 |
| | HCT | 0.060 | 0.059 | 0.044 | 0.058 | 0.026 |
| | MCH | 0.059 | 0.055 | 0.051 | 0.052 | 0.051 |
| | MCV | 0.065 | 0.064 | 0.058 | 0.056 | 0.048 |
| | ALT | 0.034 | 0.033 | 0.028 | 0.033 | 0.021 |
| | ALP | 0.099 | 0.091 | 0.079 | 0.075 | 0.060 |
| | GGT | 0.032 | 0.030 | 0.023 | 0.030 | 0.018 |
| | T2D | 0.578 | 0.619 | 0.614 | 0.549 | 0.599 |
| | BrC | 0.645 | 0.644 | 0.612 | 0.637 | 0.613 |
| | CAD | 0.594 | 0.577 | 0.564 | 0.602 | 0.558 |
| | LuC | 0.596 | 0.558 | 0.555 | 0.594 | 0.517 |

**Table S13. Real data results of five methods for $R^2$ or AUC of 26 traits in AFR, SAS, and AMR without tuning data (UKBB).**

| Pop | Trait | MIXPRS | JointPRS-auto | PRS-CSx-auto | SDPRX | XPASS |
|---|---|---|---|---|---|---|
| AFR | HDL | 0.095 | 0.090 | 0.077 | 0.084 | 0.071 |
| | LDL | 0.121 | 0.106 | 0.102 | 0.119 | 0.066 |
| | TC | 0.106 | 0.094 | 0.088 | 0.110 | 0.068 |
| | logTG | 0.050 | 0.045 | 0.036 | 0.043 | 0.034 |
| | Height | 0.095 | 0.091 | 0.062 | 0.072 | 0.053 |
| | BMI | 0.039 | 0.039 | 0.028 | 0.026 | 0.028 |
| | SBP | 0.026 | 0.016 | 0.009 | 0.023 | 0.009 |
| | DBP | 0.021 | 0.014 | 0.009 | 0.019 | 0.007 |
| | PLT | 0.050 | 0.047 | 0.038 | 0.015 | 0.021 |
| | T2D | 0.567 | 0.567 | 0.554 | 0.555 | 0.543 |
| | BrC | 0.540 | 0.551 | 0.530 | 0.532 | 0.551 |
| SAS | HDL | 0.156 | 0.133 | 0.109 | 0.123 | 0.100 |
| | LDL | 0.090 | 0.077 | 0.061 | 0.070 | 0.030 |
| | TC | 0.089 | 0.075 | 0.060 | 0.074 | 0.041 |
| | logTG | 0.122 | 0.108 | 0.091 | 0.107 | 0.086 |
| AMR | HDL | 0.139 | 0.140 | 0.125 | 0.103 | 0.063 |
| | LDL | 0.096 | 0.081 | 0.065 | 0.085 | 0.057 |
| | TC | 0.105 | 0.085 | 0.073 | 0.098 | 0.044 |
| | logTG | 0.094 | 0.078 | 0.060 | 0.069 | 0.040 |

**Table S14. Real data results of MIXPRS, PUMAS-EN, and PUMAS-EN_paper for $R^2$ of four lipids traits in EAS, AFR, and SAS UKBB without tuning data (UKBB).**

| Pop | Trait | MIXPRS | PUMAS-EN | PUMAS-EN_paper |
|---|---|---|---|---|
| EAS | HDL | 0.174 | 0.058 | 0.134 |
| | LDL | 0.095 | 0.033 | 0.051 |
| | TC | 0.095 | 0.014 | 0.073 |
| | logTG | 0.107 | 0.034 | 0.103 |
| AFR | HDL | 0.095 | 0.047 | 0.102 |
| | LDL | 0.121 | 0.085 | 0.083 |
| | TC | 0.106 | 0.070 | 0.106 |
| | logTG | 0.050 | 0.016 | 0.042 |
| SAS | HDL | 0.156 | 0.058 | 0.152 |
| | LDL | 0.090 | 0.023 | 0.051 |
| | TC | 0.089 | 0.026 | 0.047 |
| | logTG | 0.122 | 0.051 | 0.114 |

**Table S15. Real data results of eights methods evaluated by mean $R^2$ or AUC across 5-fold cross-validation for 26 traits in EAS when tuning and testing data are from the same cohort (UKBB).**

| Pop | Trait | MIXPRS | JointPRS | SDPRX | XPASS | PRS-CSx | PROSPER | MUSSEL | BridgePRS |
|---|---|---|---|---|---|---|---|---|---|
| EAS | HDL | 0.175 | 0.168 | 0.146 | 0.132 | 0.154 | 0.158 | 0.147 | 0.134 |
| | LDL | 0.096 | 0.086 | 0.092 | 0.074 | 0.081 | 0.075 | 0.043 | 0.051 |
| | TC | 0.098 | 0.104 | 0.092 | 0.069 | 0.088 | 0.096 | 0.074 | 0.071 |
| | logTG | 0.107 | 0.095 | 0.098 | 0.072 | 0.089 | 0.089 | 0.097 | 0.080 |
| | Height | 0.240 | 0.236 | 0.224 | 0.182 | 0.229 | 0.220 | 0.197 | 0.128 |
| | BMI | 0.071 | 0.089 | 0.084 | 0.063 | 0.097 | 0.092 | 0.090 | 0.064 |
| | SBP | 0.082 | 0.066 | 0.078 | 0.045 | 0.066 | 0.064 | 0.058 | 0.047 |
| | DBP | 0.081 | 0.080 | 0.075 | 0.047 | 0.077 | 0.070 | 0.069 | 0.039 |
| | PLT | 0.108 | 0.136 | 0.091 | 0.094 | 0.121 | 0.125 | 0.099 | 0.002 |
| | WBC | 0.081 | 0.071 | 0.076 | 0.045 | 0.066 | 0.070 | 0.073 | 0.011 |
| | NEU | 0.060 | 0.057 | 0.058 | 0.035 | 0.053 | 0.052 | 0.049 | 0.004 |
| | LYM | 0.091 | 0.096 | 0.088 | 0.044 | 0.091 | 0.090 | 0.087 | 0.005 |
| | MON | 0.086 | 0.077 | 0.083 | 0.049 | 0.080 | 0.078 | 0.086 | 0.018 |
| | EOS | 0.043 | 0.039 | 0.038 | 0.024 | 0.038 | 0.040 | 0.043 | 0.011 |
| | RBC | 0.091 | 0.088 | 0.093 | 0.066 | 0.088 | 0.092 | 0.080 | 0.005 |
| | HB | 0.055 | 0.051 | 0.055 | 0.024 | 0.055 | 0.055 | 0.045 | 0.017 |
| | HCT | 0.061 | 0.061 | 0.058 | 0.028 | 0.061 | 0.059 | 0.056 | 0.011 |
| | MCH | 0.061 | 0.061 | 0.053 | 0.053 | 0.064 | 0.063 | 0.051 | 0.008 |
| | MCV | 0.069 | 0.067 | 0.058 | 0.052 | 0.068 | 0.070 | 0.059 | 0.005 |
| | ALT | 0.042 | 0.044 | 0.041 | 0.028 | 0.045 | 0.037 | 0.034 | 0.022 |
| | ALP | 0.100 | 0.104 | 0.078 | 0.061 | 0.107 | 0.099 | 0.088 | 0.075 |
| | GGT | 0.044 | 0.051 | 0.041 | 0.024 | 0.053 | 0.054 | 0.035 | 0.048 |
| | T2D | 0.573 | 0.613 | 0.547 | 0.596 | 0.646 | 0.635 | 0.628 | 0.583 |
| | BrC | 0.638 | 0.631 | 0.635 | 0.614 | 0.568 | 0.615 | 0.612 | 0.639 |
| | CAD | 0.590 | 0.578 | 0.568 | 0.576 | 0.587 | 0.564 | 0.559 | 0.515 |
| | LuC | 0.653 | 0.644 | 0.623 | 0.575 | 0.659 | 0.284 | 0.591 | 0.592 |

**Table S16. Real data results of eights methods evaluated by mean $R^2$ or AUC across 5-fold cross-validation for 26 traits in AFR, SAS, and AMR when tuning and testing data are from the same cohort (UKBB).**

| Pop | Trait | MIXPRS | JointPRS | SDPRX | XPASS | PRS-CSx | PROSPER | MUSSEL | BridgePRS |
|---|---|---|---|---|---|---|---|---|---|
| AFR | HDL | 0.095 | 0.093 | 0.087 | 0.057 | 0.091 | 0.100 | 0.074 | 0.068 |
| | LDL | 0.122 | 0.110 | 0.124 | 0.059 | 0.113 | 0.098 | 0.078 | 0.073 |
| | TC | 0.106 | 0.100 | 0.112 | 0.052 | 0.098 | 0.100 | 0.079 | 0.069 |
| | logTG | 0.050 | 0.051 | 0.052 | 0.032 | 0.020 | 0.039 | 0.015 | 0.031 |
| | Height | 0.096 | 0.098 | 0.076 | 0.049 | 0.087 | 0.097 | 0.085 | 0.015 |
| | BMI | 0.040 | 0.039 | 0.027 | 0.028 | 0.041 | 0.043 | 0.044 | 0.027 |
| | SBP | 0.027 | 0.026 | 0.028 | 0.011 | 0.026 | 0.024 | 0.017 | 0.008 |
| | DBP | 0.021 | 0.020 | 0.020 | 0.007 | 0.020 | 0.015 | 0.014 | 0.005 |
| | PLT | 0.050 | 0.059 | 0.019 | 0.025 | 0.057 | 0.061 | 0.049 | 0.028 |
| | T2D | 0.567 | 0.568 | 0.556 | 0.545 | 0.558 | 0.561 | 0.564 | 0.537 |
| | BrC | 0.542 | 0.553 | 0.544 | 0.555 | 0.586 | 0.575 | 0.576 | 0.574 |
| SAS | HDL | 0.155 | 0.149 | 0.124 | 0.084 | 0.144 | 0.158 | 0.130 | 0.129 |
| | LDL | 0.089 | 0.084 | 0.070 | 0.034 | 0.081 | 0.093 | 0.085 | 0.056 |
| | TC | 0.088 | 0.087 | 0.073 | 0.039 | 0.084 | 0.085 | 0.092 | 0.064 |
| | logTG | 0.123 | 0.113 | 0.108 | 0.080 | 0.114 | 0.117 | 0.101 | 0.096 |
| AMR | HDL | 0.155 | 0.162 | 0.120 | 0.068 | 0.154 | 0.154 | 0.133 | 0.097 |
| | LDL | 0.098 | 0.089 | 0.088 | 0.060 | 0.056 | 0.058 | 0.090 | 0.043 |
| | TC | 0.121 | 0.097 | 0.121 | 0.066 | 0.093 | 0.113 | 0.098 | 0.072 |
| | logTG | 0.084 | 0.090 | 0.060 | 0.046 | 0.073 | 0.077 | 0.064 | 0.081 |

**Table S17. Real data results of eights methods evaluated by $R^2$ for nine traits in AFR and AMR when tuning and testing data are from different cohorts (UKBB and AoU).**

| Pop | Trait | MIXPRS | JointPRS | SDPRX | XPASS | PRS-CSx | PROSPER | MUSSEL | BridgePRS |
|---|---|---|---|---|---|---|---|---|---|
| AFR | HDL | 0.072 | 0.063 | 0.069 | 0.057 | 0.063 | 0.071 | 0.053 | 0.059 |
| | LDL | 0.065 | 0.059 | 0.065 | 0.036 | 0.058 | 0.050 | 0.044 | 0.046 |
| | TC | 0.055 | 0.046 | 0.052 | 0.032 | 0.049 | 0.050 | 0.047 | 0.037 |
| | logTG | 0.054 | 0.048 | 0.049 | 0.042 | 0.032 | 0.037 | 0.036 | 0.025 |
| | Height | 0.094 | 0.099 | 0.085 | 0.054 | 0.094 | 0.093 | 0.085 | 0.021 |
| | BMI | 0.030 | 0.028 | 0.025 | 0.020 | 0.030 | 0.031 | 0.029 | 0.020 |
| | SBP | 0.009 | 0.011 | 0.010 | 0.005 | 0.011 | 0.009 | 0.008 | 0.003 |
| | DBP | 0.008 | 0.010 | 0.009 | 0.005 | 0.010 | 0.008 | 0.009 | 0.004 |
| | PLT | 0.050 | 0.055 | 0.016 | 0.031 | 0.048 | 0.052 | 0.052 | 0.028 |
| AMR | HDL | 0.144 | 0.139 | 0.107 | 0.059 | 0.132 | 0.137 | 0.133 | 0.112 |
| | LDL | 0.069 | 0.067 | 0.053 | 0.016 | 0.062 | 0.059 | 0.055 | 0.028 |
| | TC | 0.078 | 0.076 | 0.070 | 0.032 | 0.068 | 0.069 | 0.071 | 0.049 |
| | logTG | 0.126 | 0.110 | 0.102 | 0.060 | 0.093 | 0.110 | 0.031 | 0.092 |

**Table S18. Real data results of MIXPRS, IndPRS, and SDPRX evaluated by $R^2$ for nine traits in AFR and AMR when tuning and testing data are from different cohorts (UKBB and AoU).**

| Pop | Trait | MIXPRS | IndPRS | SDPRX |
|---|---|---|---|---|
| AFR | HDL | 0.072 | 0.072 | 0.069 |
| | LDL | 0.065 | 0.068 | 0.065 |
| | TC | 0.055 | 0.057 | 0.052 |
| | logTG | 0.054 | 0.052 | 0.049 |
| | Height | 0.094 | 0.102 | 0.085 |
| | BMI | 0.030 | 0.031 | 0.025 |
| | SBP | 0.009 | 0.012 | 0.010 |
| | DBP | 0.008 | 0.011 | 0.009 |
| | PLT | 0.050 | 0.059 | 0.016 |
| AMR | HDL | 0.144 | 0.152 | 0.107 |
| | LDL | 0.069 | 0.069 | 0.053 |
| | TC | 0.078 | 0.074 | 0.070 |
| | logTG | 0.126 | 0.124 | 0.102 |
